# Patient-derived zebrafish xenograft models reveal ferroptosis as a fatal and druggable weakness in metastatic uveal melanoma

**DOI:** 10.1101/2021.10.26.465874

**Authors:** Arwin Groenewoud, Jie Yin, Maria Chiara Gelmi, Samar Alsafadi, Fariba Nemati, Didier Decaudin, Sergio Roman-Roman, Helen Kalirai, Sarah E. Coupland, Aart G. Jochemsen, Martine J. Jager, Ewa Snaar-Jagalska

## Abstract

Uveal melanoma (UM) is the most common intraocular melanoma, derived from transformed melanocytes of the uvea. Although treatment of primary UM is usually successful, there is a high risk (up to 50%) of liver metastasis with negligible long-term survival. There are currently no reproducible patient-derived animal models that faithfully recapitulate the latter stages of metastatic dissemination of UM, hindering the discovery of curative treatments. To overcome this problem and to accelerate the development of new metastatic UM treatments, we developed a patient-derived zebrafish xenograft (zf-PDX) model, using spheroid cultures generated from metastatic and primary UM tissues. Engrafted UM cells derived from these spheroid cultures give rise to metastatic lesions and recapitulate the molecular features of UMs and their potential drug sensitivity. Importantly, harnessing this versatile model, we reveal a high sensitivity of circulating UM cells to ferroptosis induction *in vivo* by Erastin and RSL3. Our findings are further corroborated by supportive analysis of patient data implicating ferroptosis as a new, and druggable, target for the treatment of metastatic UM patients, specifically in those with BAP1 loss in the tumor.

## Introduction

Uveal melanoma (UM) is an aggressive and deadly ocular cancer, derived from melanocytic cells of the uvea (made up of the iris, choroid, and ciliary body). UM usually carry a low mutational burden when compared to other melanomas. Strikingly, UM almost obligately bear an inactivating *GNA* family mutation (mainly in *GNAQ* and *GNA11*), blocking GTPase activity within this catalytic subunit of the protein, effectively driving oncogenic hyperactivation of G_q_ or G_11_ . This hyperactivation leads to a subsequent increase in downstream signaling, including the protein kinase C (PKC)/ MAP kinase/ ERK axis^2, 3^. UM is characterized by strong prognosticators such as monosomy 3^4–8^ and the loss of expression of the *BRCA*-associated-protein 1 (*BAP1*) gene located on chromosome 3, which is usually accompanied by the loss of chromosome 3^9^. Between 7-33% of all primary UM patients develop deadly metastatic disease within 10 years, and this is strongly linked to mutations in the *BAP1* gene. Primary UM is commonly treated by radiotherapy or by enucleation (surgical removal of the eye)^10^. Although this generally leads to effective local control, the prognosis of metastatic UM patients is grim, with a median survival of 3.9 months after detection of metastases^11^. Metastatic UM responds poorly to conventional and targeted chemotherapy^12^. In contrast to cutaneous melanomas, UM is largely refractory to immunotherapy, probably due to its low mutational burden^13, 14^.

Although metastatic spread of cancer kills the vast majority of cancer patients, the process in itself is vastly inefficient, with between 90-99% of all circulating cancer cells dying before finding a suitable metastatic niche^15–17^. This fatal weakness of UM cells limits dissemination and has hampered successful generation of animal models for therapy development. Conversely, this does highlight an exploitable opportunity for the development of novel treatments. Recent discoveries have uncovered the role of ferroptosis in the suppression of metastasis development, contributing to the attrition of circulating tumor cells^18, 19^.

Ferroptosis is a non-apoptotic form of regulated cell death that is caused by cystine depletion and overproduction of lipid-based reactive oxygen species (ROS), particularly lipid hydroperoxide, in an iron-dependent manner. SLC7A11, the catalytic subunit of the cystine/glutamate antiporter (system X_c_−), is the major transporter of extracellular cystine. Intracellular cystine is rapidly converted to cysteine and serves as the precursor for glutathione synthesis. Glutathione peroxidase 4 (GPX4) protects cells against membrane lipid peroxidation and inhibits ferroptosis. In brief, *GPX4* enzymatically reduces oxidized phospholipids, under the presence of intracellular glutathione. Broadly speaking, either inhibition of *GPX4*, or lowering of the amount of available glutathione, would enhance the levels of ROS, thereby inducing ferroptosis ^19^. Cells that exhibit oncogenic hyperactivation of the RAS-signaling cascade are sensitized to this type of cell death due to a purported de-regulation of iron homeostatic mechanisms concurrently affected^20^. The fact that over 90% of UM carry somatic *GNAQ/*11 mutations^21^ which are known to activate the RAS-MAP kinase pathway, and SLC7A11 being reported as a key downstream target of BAP1^22^, prompted us to consider ferroptosis as a druggable pathway in UM. Paradoxically, in contrast to the lethal nature of metastatic UM, no reproducible metastatic xenograft models have been established that are suitable for drug screening. Recently generated, patient-derived murine xenograft (PDX) models have been used to screen new drug combinations^23–25;^ however, the PDXs are commonly grown subcutaneously and do not resemble dissemination of metastatic disease. Concordantly, this lack of metastatic UM models limits the assessment of new treatment strategies. We have sought to mitigate this shortcoming through the establishment of a versatile, UM patient-derived zebrafish xenograft model (UM zf-PDX), combined with a novel 3D spheroid culture method for UM to ensure that metastatic properties are maintained, in order to recapitulate the final stages of the metastatic cascade.

Here we report the generation of a spheroid-derived UM zf-PDX model, and its use for the screening of ferroptosis inducers on metastatic UM. We assess that circulating UM cells are extremely sensitive to ferroptosis induction *in vivo*. We have used this druggable weakness and determined that conventional ferroptosis activators are potent inducers of ferroptotic UM cell death in a *BAP1*-dependent manner. This new insight opens the way for possible clinical treatment with ferroptosis inducers, after *BAP1* stratification of UM patients.

## Materials and methods

### Adherent cell culture of UM cells

UM cell lines MP46, MM28, and Xmm66 were provided by Dr. Samar Alsafadi (Institute Curie)^24^ and cell lines Omm1^26^, Mel285^27^, and Omm2.3^27^ from Dr. Aart G. Jochemsen (Leiden University Medical Center), respectively. All lines were cultured in Dulbecco’s modified eagle’s medium (DMEM) containing 10% fetal bovine serum (FBS, Gibco, Thermo Fisher Scientific, Waltham, MA USA), supplemented with Glutamax (Gibco). Cell lines were kept in culture up to 20 passages and intermittently checked for the presence of mycoplasma, using the PCR-based mycoplasma detection kit from American type cell culture (ATCC, Mycoplasma detection kit), following the manufacturer’s prescriptions.

### Lentiviral transduction of UM spheroid cultures

Both adherent cell culture and spheroid cultures were lentivirally transduced as described in Heitzer *et al.,* 2019^28^. In brief, the adherent cell cultures were cultured in the presence of lentiviral particles containing ΔLTR flanked CMV:tdTomato-blasticidin (Addgene#106173) and 8µg/mL polybrene (Sigma, Zwijndrecht, the Netherlands), whereafter the medium was exchanged for standard culture medium. Transduced adherent UM lines were selected with 2µg/mL blasticidin (Gibco) for approximately 3 passages (approximately 10 days) until all cells were positive for the transduced tdTomato construct. For obtaining a cultured spheroid, the procedure was the same, except of the generation of a single cell suspension prior to the addition of the viral particles and the omission of the selection.

### Establishment of stable PDX-derived spheroid cultures

Metastatic PDX tissues that had been frozen down in either FBS containing 10% dimethyl sulfoxide (DMSO) or primary PDX tissues that were frozen in neuronal stem cell medium (NSC medium, Stemcell technologies, Köln, Germany) containing 10% DMSO were thawed by brief incubation at 37°C and were transferred to basal NSC medium. Subsequently, the medium was exchanged for 10 mL DMSO-free medium containing 5 mg/mL Primocin (Invivogen, Toulouse, France), the tissue was minced in 10mL basal NSC in a cell culture petri dish using a sterile scalpel blade. The material was collected in a 50mL centrifuge tube and supplemented with 0.01 mg/mL Liberase TL (Roche, Woerden, the Netherlands); the suspended tumor tissue was incubated at 37°C for 3-5 hours while shaking at 250 rpm, and the tubes were subsequently vortexed intermittently during this incubation to break up any tissue aggregates. The disaggregation process was monitored macroscopically; at the end of the procedure a small sample was observed under an inverted microscope to ensure completion of dissociation, otherwise the dissociation was prolonged. After the cell suspension was passed through a sterile 30 µm cell strainer to remove all cell and extra-cellular matrix aggregates, cells were pelleted and suspended in complete NSC medium (supplemented with both 1x B27 (Gibco) and 1x N2 (Gibco), 20 ng/mL bFGF (Peprotech, Hamburg, Germany), 20 ng/mL EGFP (Peprotech), 5 U/mL heparin and 1x primocin (Invivogen), containing 5% FCS and 200mM Glutamax). The cell suspension was diluted and plated in a 24-well ultra-low attachment plate (Corning, Wiesbaden, Germany), in approximately 8-12 wells with a 0.25 cm^3^ tumor volume at the start of the dissociation. After several days of culture, the cells coalesced into larger cell aggregates to be disrupted prior to labelling and engraftment.

### Staining of UM spheroid culture derived cells prior to implantation into embryonic zebrafish hosts

Spheroids were collected and concentrated through centrifugation (200 x g, 5 min). The cells were resuspended in 3mL TrypLE (Gibco), and after a 10-minute incubation at 37°C, combined with intermittent agitation with a 1000µL pipette cells, aggregates were broken up by pipetting up and down. TrypLE was subsequently inactivated by addition of 7mL complete NC medium, followed by centrifugation.

The red intra lipid dye CM-DiL (Sigma) was used to stain the cells to visualize cancer cell proliferation and metastatic initiation. In brief, the disaggregated cell suspension was concentrated to 2 mL in NC complete medium in a 15mL tube and supplemented with 2.5 µM CM-DiL followed by 30 min incubation at 37°C in the dark. The unbound labelling reagent was removed through centrifugation after addition of 8 mL complete NC medium.

### Establishment of maximum tolerated drug dose in zebrafish

Prior to either single or combinatorial drug treatment on engrafted zebrafish larvae, we established a maximum tolerated dose (MTD), where we have at least 80% survival of the treated un-injected larvae. To achieve this, we crossed Casper mutant zebrafish, raised the larvae up to 2 days post fertilization (dpf) and placed 6 larvae per well in a cell-culture grade 24-well plate (Corning). We subjected the larvae to a concentration range of 10µM-156 nM of all tested compounds, in a two-fold dilution series. All compounds were changed every other day to ensure optimal stability of drug levels throughout the duration of the treatment. At 8dpf (corresponding to 6dpi for injected larvae), we scored survival and plotted the survival using Graphpad Pro 8 (Graphpad software LLC, San Diego, CA). The highest concentration at which over 80% of the treated larvae survived was chosen as the MTD for the treatment of the engrafted larvae. For combinatorial treatments to determine synergism, we followed the titrated MTD of compound A (the purported sensitizer) with a similar dilution series of compound B (the purported synergistic compound). We titrated from MTD A combined with 10µM-156 nM compound B to attain a suitable treatment concentration where >80% of all treated larvae survived up to 8dpf for a 6-day treatment (starting at 2dpf).

### Preparation of cells for implantation into embryonic zebrafish host

Cells were prepared for engraftment in accordance with the protocol published by Groenewoud *et al.,* 2021^29^. To facilitate the engraftment of single cells, cells were disaggregated immediately prior to implantation, concentrated by centrifugation at 200 x g for 5 minutes, after which the supernatant was removed and the pellet resuspended in 3 mL TrypLE (Gibco). This was followed by a 10-minute incubation with intermittent agitation (gentle vortexing every 2 minutes). The proteolytic activity of TrypLE was negated through the addition of 7 mL spheroid culture medium after which the cell suspension was pelleted at 200 x g. The cells were washed with Dulbecco’s PBS without Ca^2+^ and Mg^2+^ (DPBS, Gibco). PBS was removed after 5 minutes of centrifugation at 200 x g and subsequently after 30 seconds at 200 x g to ensure that all remaining DPBS is removed and can completely be replaced with sterile 2% PVP_40_ in DPBS. Cells were injected at a concentration of 250 x 10^6^ cells/mL. The cells were transferred into a glass capillary needle (needle preparation as described in Groenewoud *et al.* 2021^29^), using a micro loader tip (Eppendorf, Nijmegen, the Netherlands).

### Injection of cancer cells into zebrafish

Either *tg(fli:GFPx casper)*^30, 31^ or *tg(casper)*^32^ fish were crossed prior to the start of the experiment, and larvae were cleaned every day after harvesting up to 2dpf. The larvae were collected after they had hatched from the chorion, and the water was removed along with the chorion debris, together with all unhatched larvae (unhatched larvae were removed with the same strainer as was used to collect the eggs at harvesting).

Approximately 300-400 cells were injected into the duct of Cuvier (doC, the embryonic common cardinal vein) of 2dpf zebrafish larvae. After injection, dead larvae were removed and the residual larvae were placed in clean eggwater. The larvae were screened using a fluorescent stereo microscope, selecting all individuals that displayed no bodily malformation and that had clearly visible cell accumulation in the caudal vein and caudal hematopoietic tissue. When using Casper transgenic zebrafish, solely the presence of cells in the tail and lack of malformations was used as a screening criterion. All positively-selected individuals were moved to a clean petri dish, and after completing the screening, the positively-selected pool of individuals was screened once again to ensure that no un-injected or otherwise aberrant individuals were placed in the treatment pool.

### Confocal imaging of zebrafish xenografts

Zebrafish were anaesthetized with 0.002% tricaine (MS222, Sigma) in eggwater and embedded in 1% low melting temperature agarose dissolved in eggwater. The larvae were positioned with a trimmed down microloader tip (Eppendorf) as to be laterally oriented, gently pressing the larvae down to ensure close proximity to the lens of the confocal microscope and a level orientation of the larvae. Images were captured of both green (GFP) and red tdTomato/CMDiI channels and were recorded as approximately 1 x 4 stitches at 10x magnification using a Leica sp8 confocal microscope (Leica, Wetzlar, Germany). Consecutive stitch sequences were processed into a single image using Fiji ^33^ using the plugin by Preibisch et al., 2009 ^34^.

### IHC analysis of engrafted zebrafish larvae

Engrafted zebrafish were euthanized with tricaine and fixed for 16 hours in ice cold 4% paraformaldehyde in PBS. After fixation, larvae were washed with PBS containing 0.05% tween 20 (v/v) and 200mM Glycine. Larvae were stored in the dark at 4 °C until further processing. Fixed zebrafish larvae were arrayed in a grid and embedded in agarose (sphereoQ, Hispanagar, Burgos, Spain). Care was taken to ensure equal localization in the *x, y*, and *z* axes. Larvae were sectioned along the ventral axis, taking care to section through the tailfin and caudal hematopoietic tissue.

Sections were cut at 4 µm from formalin-fixed paraffin embedded (FFPE) blocks of UM cells-containing zebrafish as detailed above, and placed onto X-tra adhesive slides (Leica Biosystems, Milton Keynes, UK). Immunohistochemical (IHC) staining was performed using the Bond RXm Automated Stainer with high pH antigen retrieval and the Bond polymer-refine detection systems in either red or brown chromogen, according to the manufacturers’ recommendations (Leica Biosystems). Primary antibodies included mouse anti-melanA (Dako, Agilent, Cheshire UK) and mouse anti-BAP1 (Santa Cruz Biotechnology, USA), both at a concentration of 1 µg/mL. Slides were counterstained with haematoxylin and mounted with a resin-based mountant. Human UM tissue was used as a positive control for each of the primary antibodies. Mouse IgG1 isotype control at a concentration of 1 µg/mL was also included in each assay.

### Drug treatment of engrafted UM zf-PDX

All drugs were acquired from Cayman chemical (Ann Arbor, Michigan, USA) and were dissolved in dimethyl sulfoxide (DMSO) unless otherwise stated. All drugs were added at the beginning of the experiment, right after screening of the larvae (in the morning after injection at 1dpi). The drug-containing eggwater was exchanged every other day.

After careful screening of the engrafted zebrafish, larvae were randomly subdivided into a 24-well plate, with 6 individuals per well and 6 wells per condition. After plating, the eggwater was gently removed, without disturbing the larvae. Subsequently, the compounds were added, dissolved in eggwater. The volume of vehicle control used was the same as the highest volume of drug added to the plate. Subdividing the larvae in this manner allowed for the screening of 3 mono treatments combined with one vehicle control or one set of drug combinations, with vehicle control, compound A, compound B and compound A+B.

At 6dpi, the larvae were pooled per condition into a 6-well plate (Corning) and as much as possible of the drug-laced eggwater was removed; after this, the larvae were washed 3 times with 5 mL eggwater to remove all traces on non-internalized drug. From this pool of larvae, 20 random individuals were selected and imaged using a MZ16FA fluorescence microscope equipped with a DFC420C camera (Leica, Wetzlar, Germany). The microscope was set (exposure time and gain) on the control group of each experiment to ensure that there was no signal saturation in the control group and that all larvae with reduced tumor burden would fall within the set margins; focus was adjusted per larva when required. All remaining groups were imaged using the same settings.

### Clinical data analysis

The LUMC cohort includes clinical, histopathological, and genetic information on 64 UM cases enucleated between 1999 and 2008 at the Leiden University Medical Centre (LUMC). Clinical information was collected from the Integral Cancer Center West patient records and updated in 2019. For each sample, part of the tumor was snap frozen with 2-methyl butane and used for mRNA and DNA isolation, while the remainder was embedded in paraffin after 48 hours of fixation in 4% neutral-buffered formalin and was sent for histological analysis. Chromosome status was determined with the Affymetrix 250K_NSP-chip and Affymetrix Cytoscan HD chip (Affymetrix, Santa Clara, California, United States of America). RNA was isolated with the RNeasy mini kit (Qiagen, Venlo, The Netherlands) and mRNA expression was determined with the HT-12 v4 chip (Illumina, San Diego, California, United States of America).

Statistical analyses of the LUMC cohort were carried out in SPSS, version 25 (IBM Corp). For survival analysis, Kaplan-Meier and log-rank test were performed with death due to metastases as endpoint. Cases that died of another or unknown cause were censored. The two subpopulations that were compared in each analysis were determined by splitting the total cohort along the median value of mRNA expression for each analyzed gene.

### Zebrafish data acquisition and statistical analysis

All zebrafish larval engraftments were performed in a biological duplicate, unless otherwise stated, with >20 individuals per group per biological repeat. All larvae were randomized and entered into either control or experimental groups. For imaging, larvae were randomly selected and imaged using the same exposure setting with a fluorescent stereo microscope. Outliers were removed from all data sets using Graphpad Prism 8.0, (Q5) prior to normalization and combination of all biological replicates. Data were normalized to either control (drug treatment) or to day one (in growth kinetic experiments). Statistical significance was tested with an ANOVA for normally distributed data sets, while otherwise a Kruskal-Wallis test was used. Error bars depict ±SEM. Data are presented as mean ±SEM or mean ±SD. P-values ≤0.05 are considered to be statistically significant (*p 0.05, **p < 0.01, ***p < 0.001, ****p< 0.0001).

### *In vitro* growth assay

To investigate the effect(s) of inducers (Erastin and RSL3) and inhibitors of ferroptosis (Ferrostatin-1 and Liproxstatin) on cell survival *in vitro,* cell lines were seeded in triplicate or quadruplicate in 96-well plates. The next day, cells were treated with the different compounds. Survival was determined after 5 days of incubation using the Cell Titre-Blue assay (Promega). All cell lines were treated with 4 and 8 µM Erastin and 3 and 6 µM RSL3, with the exception of Mel285 which was treated with 0.05 and 0.2 µM Erastin or RSL3.

### qPCR analysis

Spheroid cells were harvested (1x10^6^) by centrifugation (200 x g for 5 min at 25°C), or in case of adherent cells, after prior trypsinization. Whole RNA was isolated using the Qiagen RNeasy kit (Qiagen) according to the manufacturer’s description, reducing sample viscosity by passing the cell lysate 5 times through a sterile 20-gauge needle and treating the isolate on-column with RNase free DNAse (provided by the manufacturer) for 15 minutes at room temperature. Total RNA yield was quantified using Nanodrop 2000 measurement (Thermo Scientific, Wilmington, USA) and cDNA was synthesized using the iSCRIPT cDNA kit (Biorad, Hercules, USA) according the manufacturer’s description, to a total of 1 µg for each cell line.

Detection was performed using the iQ5 QPCR apparatus (Biorad), using IQ green super mix (Biorad), for 35 cycles, followed by a high-resolution melting curve. All primers (supplementary table ST2, with the exception of OCT3/4 and Nanog which were taken from Chen et al 2017^35^) were diluted in PCR grade nuclease-free water (Gibco) at a concentration of 100 µM. All primers passed an efficiency test prior to use at a final concentration of 10 pmol.

Glyceraldehyde-3-phosphate dehydrogenase (GAPDH) and Calpain Small Subunit 1 (CSPN1) levels were used as an internal reference for each experimental primer set. Transcript levels were determined using the ΔCT method (when determining transcription levels without second internal normalizer, i.e., GPX4 levels in correlation with *BAP1* status) or ΔΔ CT when using an internal reference (i.e., comparison of adherent and suspended cells).

### Protein lysates and Western blot

To determine protein expression, cells were seeded into 6-well plates. After two days, when cells were ∼70-80% confluent, they were rinsed twice with ice-cold PBS on ice and subsequently lysed in Giordano buffer (50 mM Tris-HCl at pH 7.4, 250 mM NaCl, 0.1% Triton X-100, and 5 mM EDTA; supplemented with protease- and phosphatase inhibitors) for 10 minutes on ice. After scraping and transferring lysates to tubes, lysates were centrifuged for 15 minutes at 15,000 rpm. Supernatant was transferred to a clean tube.

Lysates of primary UM samples were also made in Giordano buffer, after crushing nitrogen-frozen pieces of tumor to powder, and further processed as described for the cell lines. Subsequently, protein concentrations were determined with Bradford reagent (Bio-Rad). Equal amounts of proteins were separated on SDS-polyacrylamide gels, and proteins were transferred onto PVDF membranes (Millipore). After blocking in 10% non-fat dry milk in TBST (10 mM Tris-HCl, pH 8.0, 150 mM NaCl, 0.2% Tween-20), the blots were incubated overnight with appropriate antibodies diluted in TBST, 5% BSA. After washing with TBST and incubation with appropriate secondary antibodies coupled to HRP for 30 minutes, blots were washed thoroughly and imaged using a Chemidoc (Bio-Rad). The following antibodies were used: anti-*GPX4* (B12) and anti-BAP1 (C4) (Santa Cruz Biotechnology), anti-SLC7A11 and anti-ERK1/2 (Cell Signaling Technology), anti-di-phospho-ERK and anti-Vinculin (Sigma-Aldrich).

## Results

### Metastatic capacity of Xmm66 spheroid-derived cells is lost upon adaptation to adherent culture

During development of a zebrafish metastatic UM xenograft model for drug screening, we observed a low engraftment rate of adherent UM cell lines (Figure 1A) which hindered use of this model for robust drug screening. To address this issue, we compared the *in vivo* behavior of cells generated from spXmm66 spheroids with the adherent Xmm66 cell line derived from the same metastatic tumor tissue, and with another adherent UM cell line, Omm2.3^24, 27^. After intravenous injection into zebrafish embryos, we indeed observed a significant (p<0.001) enhancement of the tumor cell burden induced by engraftment of cells derived from spXmm66 spheroids (Figure 1A) when compared to both adherent cell lines.

**Figure 1.**
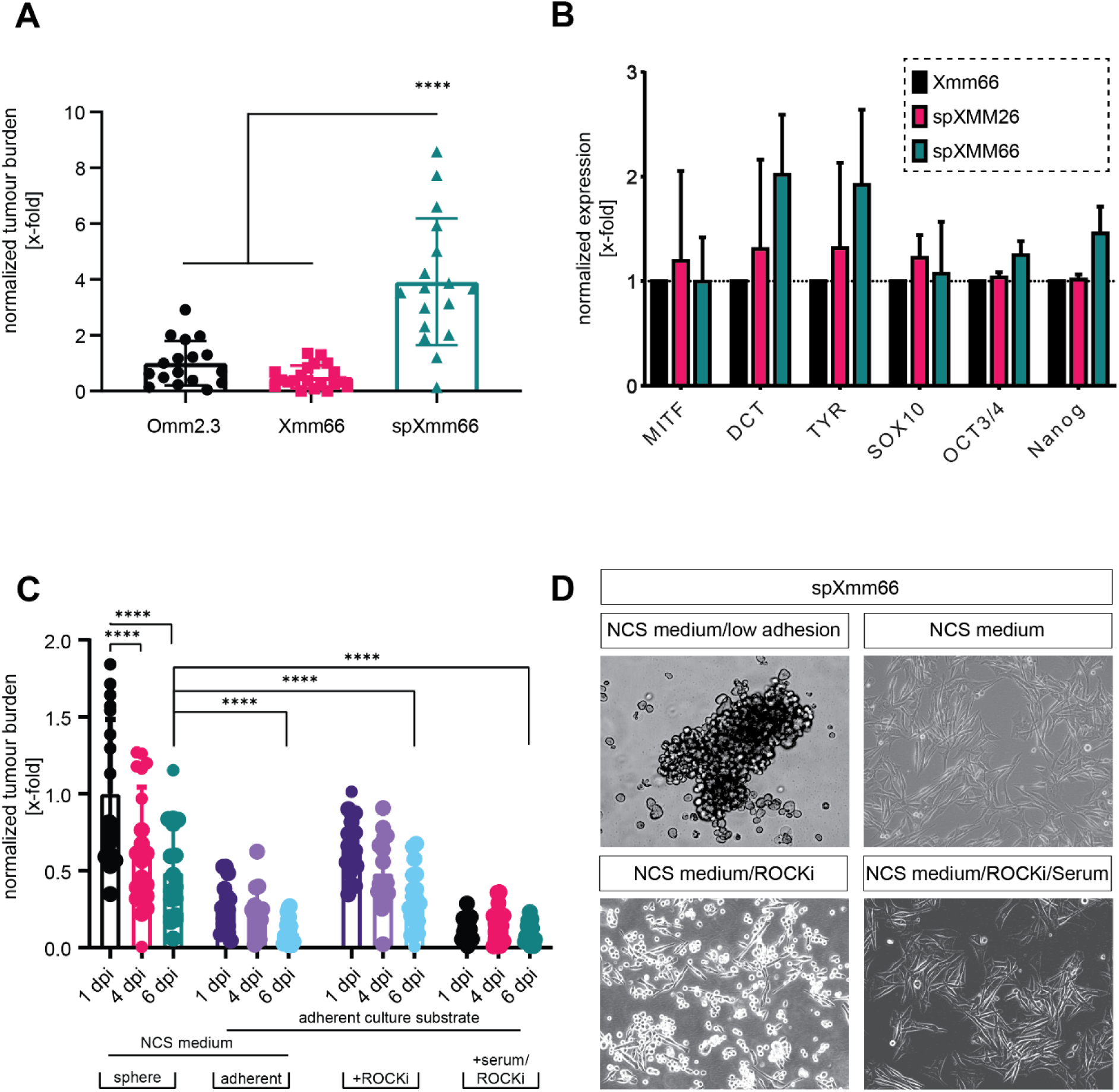
Loss of metastatic capacity of Xmm66-derived cancer cells upon shift to adherent culture. A) Metastatic capacity of adherent uveal melanoma cells and UM-derived spheroid line spXmm66 in zebrafish, n=20, error bar represents ±SEM. B) Transcriptional analysis of spheroidal uveal melanoma cell lines compared to adherent uveal melanoma cell line Xmm66, expression levels normalized to Xmm66. C) Comparison of zebrafish tumor burden after injection of near patient spheroid line spXmm66, compared to *de novo* adherent cultures, derived from spXmm66, cultured as a conventional cell culture on plastic (7 days), with the addition of Rock inhibitor Y276321 or the addition of both Rock inhibitor and fetal calf serum; each group n=20, error bars represent ±SEM. D) Microscopic images of the spXmm66 spheroid line when in suspension (on ultra-low adhesion plastic, in neuronal stem cell medium), in NSC medium on conventional cell culture plastic, in NSC medium containing ROCK inhibitor Y27632 and in NSC medium containing ROCK inhibitor Y27632 with 10% FCS.

Next, we asked whether this difference could be due to an overall loss of tumorigenic capacity, or a loss of stem cell-like features under adherent conditions, which is retained in spheroid cultures. Therefore, we compared the transcriptional activity of two spheroid cultures (spXmm26 and spXmm66) with the adherent cell line Xmm66. We found clear, yet statistically insignificant enhancements in the spheroid cultures, for melanocyte differentiation markers microphtalmia-associated transcription factor (MITF), SRY-related homology box (SOX10), dopachrome tautomerase (DCT) and tyrosinase (TYR). Furthermore, we observed an overall enhancement of “Yamanaka factors” OCT3/4 and Nanog specifically in the spXmm66 spheroid culture and not in spXmm26 or adherent Xmm66 (Figure 1B),^36^ indicating an overall enhancement of both differentiated melanocyte markers as well as stem-like markers, indicative of the presence of several discrete differentiation states, or a heterogeneous cell population within our spheroid cultures.

To test whether an inherent loss of metastatic potential of UM cells occurs upon adherent cultivation, we derived *de novo* adherent cell lines (spXmm66-Adherent) from spheroid culture spXmm66 using different growth conditions. These novel adherent lines were generated by seeding spheroid cultured cells in the neuronal stem cell (NSC) medium in conventional cell culture flasks and transducing them with the same lentiviral construct, CMV:tdTomato-blasticidin. We controlled for the effect of the NCS medium by generating one line in the presence of basal NCS medium. Subsequently, we grew two other lines in NCS medium, with or without 10% FCS. After the establishment of these *de novo* adherent cell lines, cells were implanted into the doC of 48hpf zebrafish larvae and imaged at 1, 4, and 6dpi. We observed that cells derived from spXmm66 lost their metastatic potential after the shift to the 2D substrate through adhesion. We subsequently noted that the strong negative correlation between the adhesion of cells and the metastatic potential in zebrafish could be further intensified by the addition of serum, possibly due to the pro-differentiating function of the soluble factors commonly found in FCS (Figure 1C). We subsequently assayed the effect of ROCK inhibitor Y27632 to assess the effect of ROCK-dependent signaling on the loss of metastatic potential as this inhibitor is commonly used in the organoid culture to prevent differentiation of human induced pluripotent stem cells, presumably via perturbing biomechanical signal transduction through the actin cytoskeleton^37^ (Figure 1C and D). Addition of ROCK inhibitor Y27632 to the *de novo* derived adherent cell lines inhibited the reduction of metastatic potential when compared to the suspension culture (p= 0.43), while addition of serum to the ROCK inhibitor-treated cells reduced their metastatic capacity, though not significantly (p=0.18). These findings lead us to conclude that, in UM, tumorigenic capacity is lost when cells are cultured in adherent conditions, while spheroid cultures retain the tumorigenic capacity.

### Generation of spheroid cultures derived from primary and metastatic UM tissues

To overcome the loss of metastatic capacity of UM cells upon adherent culturing we developed a spheroid culture protocol to stabilize fresh patient-derived UM material prior to further examination of drug susceptibility in the zebrafish xenograft model.

All primary tumor-derived tissue samples (n=10, representative images show samples spUM-LB046, spUM-LB048, spUM-LB049) readily formed spheroids in culture (100% success rate) within 24 hours and were cultured for 3-7 days (Figure 2A). In addition, 14 metastatic PDXs tissues were tested. One stable spheroid line (spXmm66) was generated out of 14 PDX samples (7% success rate) (Table ST1). The other 13 PDXs-derived spheroid cultures were successfully maintained as short-lived spheroid cultures for the duration of the experimental procedure (at least 7 days, 100% success). Samples derived from metastatic UM PDXs (spXmm26, 33, 66, 300) were cultured for approximately 2 months (Figure 2B). Although all primary (n=10) and metastatic (n=13) samples effortlessly formed spheroids, there was little proliferation in these samples (Table ST1), with the exception of spXmm66. The short and long-lived culture spXmm66 stained positively for melanocyte-specific antigen melanA at passages 4 and 20, indicating the retention of its melanocytic background (Figure 2C).

**Figure 2.**
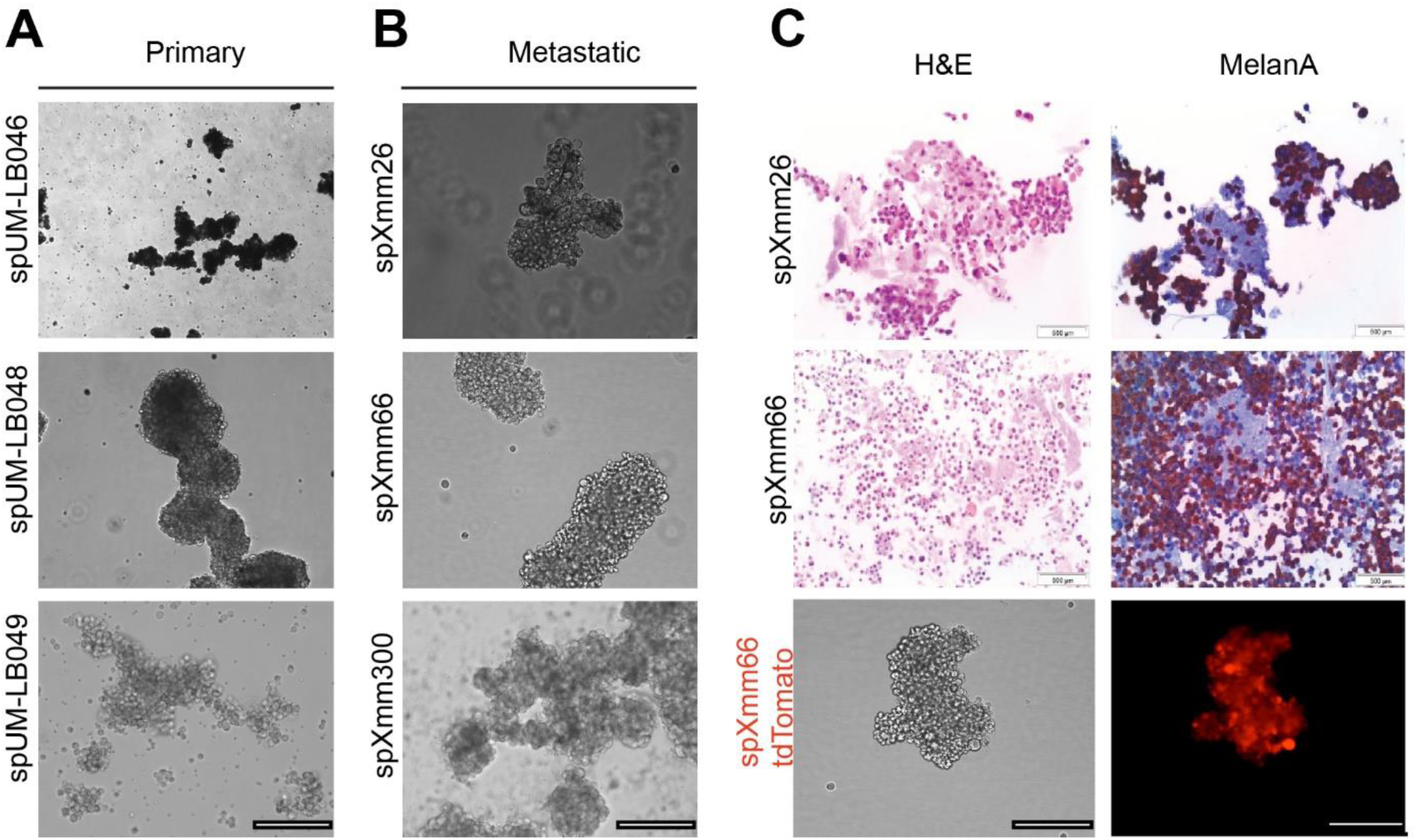
Spheroid cultures can readily be established from both primary uveal melanoma tumor and metastatic uveal melanoma PDX tissues. Representative images of the established spheroid cultures, derived from A) primary UM (spUM-LB046, spUM-LB048, spUM-LB049) and B) from metastatic murine PDX (spXmm26, spXmm66, spXmm300) (10x magnification brightfield, scalebar depicts 250 µm). C) H&E-stained metastatic spheroid cultures (pink) and spheroid cultures stained with melanocyte-specific antibody anti-melanA (magenta). Lentiviral transduced spXmm66, driving tdTomato expression in the spheroid culture derived from murine metastatic PDX material.

Importantly, we determined that the short-lived spheroid cultures could be successfully used for *in vitro* drug treatment (Supplementary Figure S1). All tested primary tissues (between 2.5-5 mm^3^ sample size during enucleation) yielded enough material after short-lived spheroid culture for at least two zebrafish engraftments within 7-14 days after establishment. Only the long-lived spXmm66 culture propagated sufficient material for repeated zebrafish engraftments, allowing single and combinatorial drug testing. Collectively, we have established a successful platform to isolate, preserve and recover viable tissue and spheroid cultures generated from UM patient material, either murine PDX or primary-tumor derived, for subsequent engraftment and validation.

### Spheroid-derived metastatic xenografts yield a reproducible metastatic phenotype and recapitulate molecular features of UM cells in zebrafish

There are currently no animal models for metastatic uveal melanoma suitable for drug screening. To address this issue, we established a zebrafish xenograft model by intravenous injection of fluorescent UM cells derived from a short-lived primary spheroid and long-lived metastatic spXmm66 spheroid cultures. Considering the limitless availability of spXmm66 cells and our interest in developing drug screening platform for metastatic UM we continued our investigation using the spXmm66 model. Engrafted fluorescent spXmm66 cells disseminated hematogenously and formed metastatic foci (Figure 3A, B). To verify the presence of viable UM cells, zebrafish engrafted with spXmm66-derived cells were selected using a fluorescent stereo microscope and subsequently fixed at 6 days post implantation, and imaged using a confocal microscope, generating 10 x whole body stitches. Representative confocal stiches indicate primary UM zf-PDX models spUM-LB048 and spUM-LB048 (Figure 3B). Paraffin-embedded zebrafish were sectioned and stained for the melanocyte-specific marker melanA and for the presence of BAP1. All engrafted larvae showed melanA and BAP1-positive cells (Figure 3C) proving that engrafted cells maintain expression of UM histological markers after injection (additional IHC’s of zf-PDX in supplementary Figure S5).

**Figure 3.**
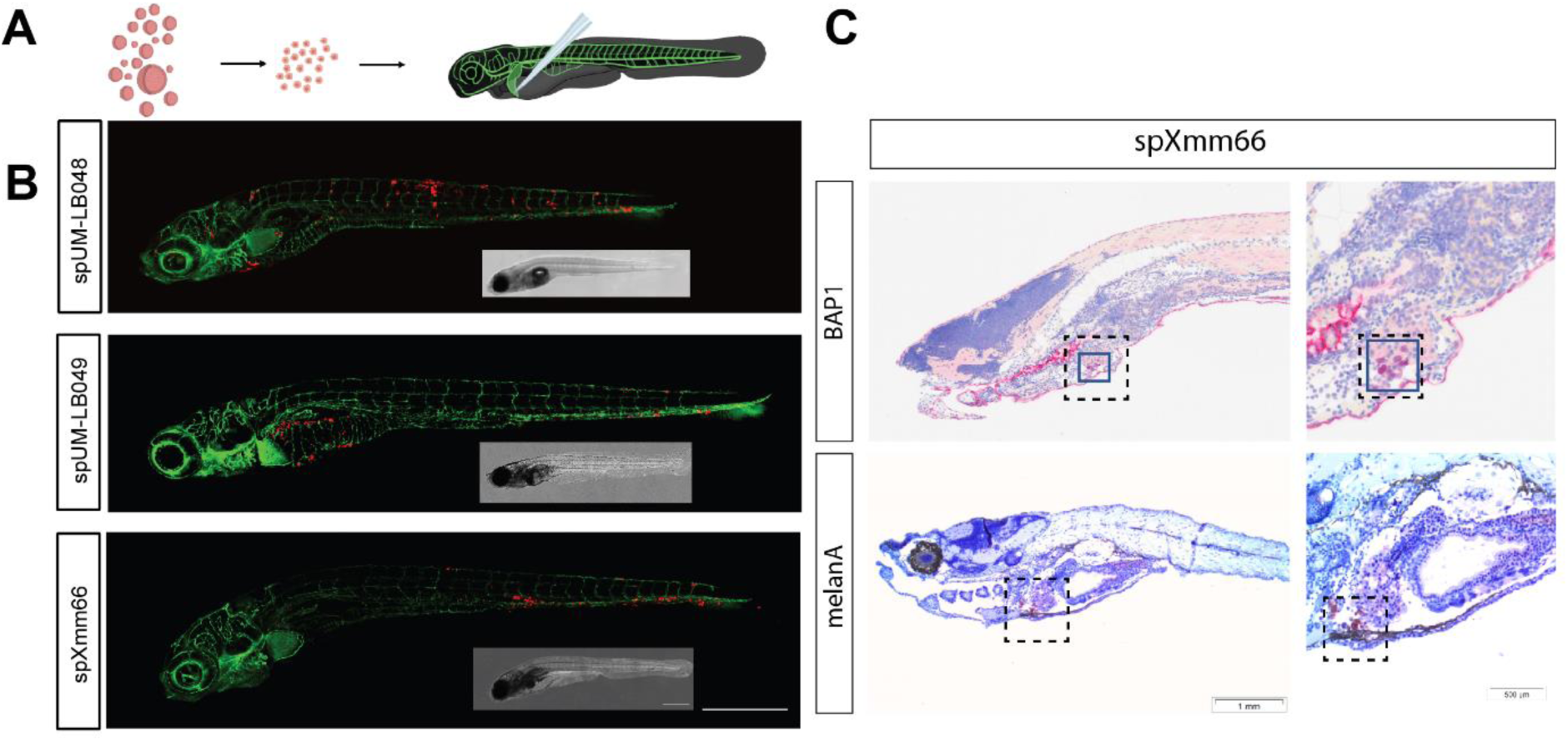
Establishment of a UM zf-PDX model through duct of Cuvier (doC) injection yields a reproducible metastatic phenotype. A) Schematic representation of the engraftment procedure; spheroid cultures are reduced to single cell suspensions by enzymatic dissociation, and single cells are injected into the blood circulation of 48 hpf Tg(fli:GFP) zebrafish larvae through the doC. B) Representative confocal micrographs of 6 days post injection, showing GFP (green) blood vessels in zebrafish, engrafted with cancer cells marked in red (either CM-DiL for spUM-LB048, 049 or tdTomato lentiviral over-expression for spXmm66); scale bar represents 250 μm. Disseminated cancer cells are present up to 6 days post engraftment and settle in both the hematopoietic tissue and the liver. C) Hematoxylin and eosin staining and BAP1 IHC (dark purple, boxed area), and melanA (dark purple, boxed area), staining on spXmm66 engrafted zebrafish larvae. Scale bar equals 1 mm and 500 µm for the magnification.

The long-lived spXmm66 spheroid culture, derived from the same metastatic tissue-derived PDX as the adherent UM cell line Xmm66 ^24^, proved to be highly proliferative and metastatic in the zebrafish xenograft model. Cancer cells were present up to 6 days post engraftment, the ethical endpoint of the experiment, with distinct cancer cell colonies arising in the liver and the caudal hematopoietic tissue of the zebrafish (Figure 3B, C).

### Combinatorial small molecule inhibitor screening validates the UM zf-PDX model as a versatile tool for anti-UM drug discovery

To confirm the validity of our metastatic UM zf-PDX spXmm66 derived model as a potential drug screening tool, we tested several known small molecule inhibitors. These therapeutics or combinatorial therapies were originally selected through a combination of genomic analysis, *in vitro* pre-screening and an efficacy study in a murine subcutaneous UM PDX model, derived from the same metastatic UM patient (Xmm66 tissue)^23^. We used spXmm66-derived cells engrafted in zebrafish and tested three combinatorial sets of small molecule inhibitors, previously published, as a means of chemical validation of the zf-PDX model^25, 38, 39^. The spXmm66 engrafted zebrafish were exposed to: mTORC1 inhibitor-everolimus (RAD001) together with PKC inhibitor-sotrastaurin (AEB071), BCL-2/BCL-xl inhibitor-navitoclax (ABT263) combined with RAD001 and HDAC inhibitor-quisinostat, and with CDK inhibitor-flavopiridol (Figure 4). Prior to treatment, we first established the maximum tolerated doses (MTD) of selected drugs by treating un-injected zebrafish larvae, from 72hpf for 5 days, changing the drug-laced zebrafish medium every other day, in a similar fashion as the final drug treatment (Figure S2). The MTD of everolimus was 2.5 µM and of sotrastaurin 2.5 µM (Figure S1). The combinations were made as previously mentioned, after subsequent titration: everolimus at 1.25 µM combined with sotrastaurin at 2.5 µM; everolimus at 1.25 µM combined with navitoclax (ABT263) at 5.0 µM; quisinostat at 500 nM with flavopiridol at 1.0 µM. We analyzed the decrease of tumor burden as described by Groenewoud *et al.* 2021^29^: all groups, either by mono treatment or combination treatment, were compared to vehicle control (Figure 4). For the combination of flavopiridol and quisinostat, we measured a significant reduction of tumor burden (p < 0.001) but not when using a single compound. For the combination of everolimus and navitoclax we saw a significant inhibition with the mono treatment of navitoclax (p<0.05), that was further enhanced by the addition of everolimus (p<0.001). After this preliminary chemical validation, we concluded that this zf-PDX model recapitulates drug sensitivity similar to corresponding murine PDX and therefore could be applied for the development of new, pre-clinical experimental treatments^24, 25, 38, 39^.

**Figure 4.**
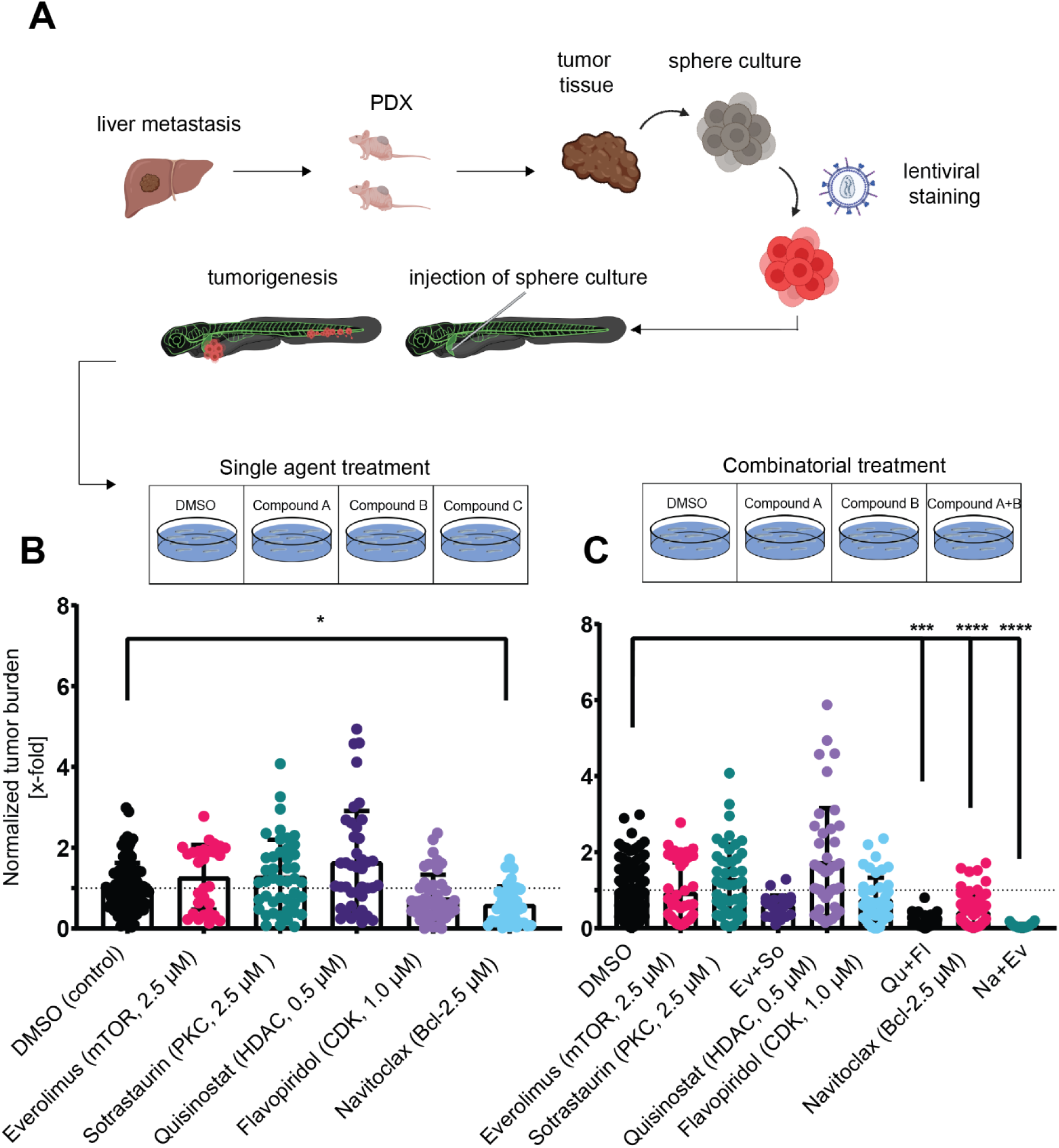
Chemical validation of the established metastatic uveal melanoma model. A) Engrafted zebrafish with UM cells derived from the spheroid cultures were treated with the maximum tolerated dose of the compounds, determined as previously described (shown in Figure S1). B, C) Tumor burden, normalized to DMSO control (normalized tumor burden); measurements are combined from at least 2 experiments and 20 individuals (n>20*2), error bar depicts ±SEM.

### Uveal melanoma cells are highly sensitive to the induction of ferroptosis *in vivo,* enabling novel anti-UM therapeutic strategies

Next, we addressed why the development of metastatic animal UM models has proven to be such a difficult challenge. During the establishment of our UM drug screening platform, we have concluded that there is a clear discrepancy between adherent cultured UM cell lines and UM cells capable of metastasis.

We hypothesized that circulating UM cells are hyper sensitive to cell extrinsic stress factors during metastatic dissemination therefore establishment of reproducible metastatic models is very inefficient process. We came to this hypothesis due to the fast and complete clearance of the engrafted adherent cells from the zebrafish host after injection (Figure 1A). We reasoned that the cells were destroyed by a cell intrinsic mechanism during metastatic dissemination. We propose that understanding of this mechanism will give us an insight into how UM cells resist clearance in circulation and induce metastasis in patient.

Recent discoveries in cutaneous melanoma metastasis and the inhibitory effects of ROS on metastasis, led us to assess the effect of some of the key regulatory proteins on UM metastasis^18, 40, 41^. Moreover, due to the fast and complete clearance of the adhered UM cells from the zebrafish host after injection (Figure 1A), we reasoned that these cells were destroyed by a cell intrinsic mechanism.

Recent developments in ferroptosis biology^42^ and the inhibitory effects of ferroptosis on disseminating cutaneous melanoma^18^ prompted us to ask if ferroptosis could be the underlying, cell intrinsic mechanism, causal to the low tumorigenic capacity and rapid clearance of UM cells. Combining these recent findings with the parallel between the rapid and complete clearance of UM cells from the engrafted zebrafish host, we hypothesized that the high metastatic potential of UM cells in patients might be explained through an upregulation of ferroptosis detoxifying mechanisms. To test this, we investigated the expression of known ferroptosis regulators and whether expression levels affect the survival of UM patients (Figure 5).

**Figure 1.**
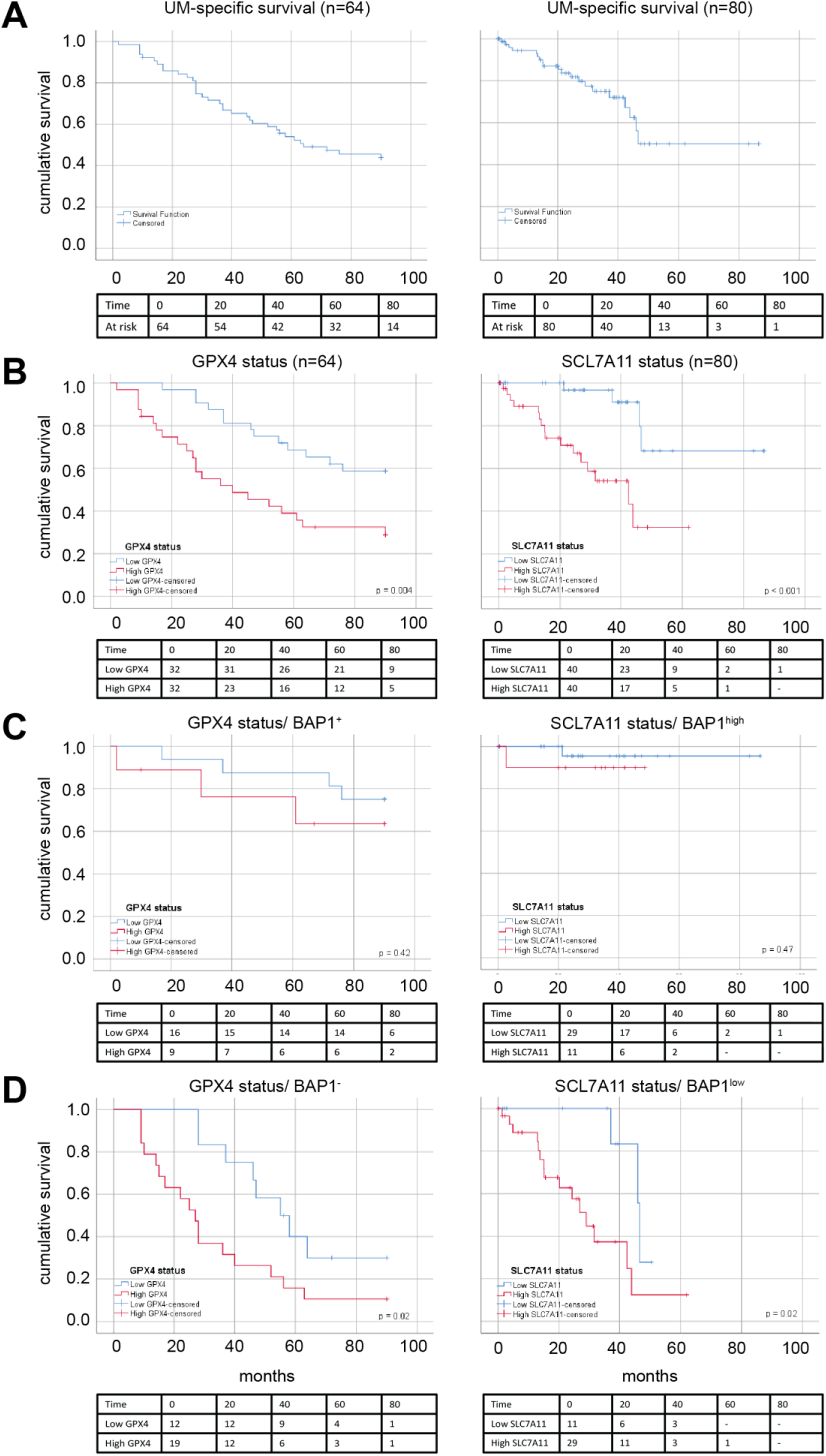
Ferroptosis detoxification mechanisms negatively correlate with UM patient survival in the Leiden cohort and with melanoma specific survival in the TCGA cohort. A) Analysis of the UM-specific survival in both LUMC UM and TCGA patient cohorts. B) UM-specific survival of GPX4 (LUMC cohort, n=64) and SCL7A11 (TCGA, n=80), expression divided over the median, shows a negative correlation of both *GPX4* (p = 0.004) and system Xc- (p = 0.0014) on patient survival C) Comparative analysis of the relation between *GPX4* and *SCL7A11* and survival in BAP1+ (LUMC, IHC, n=25) and BAP1 high (TCGA, RNAseq, n=40) UM samples and in D) BAP1 – (LUMC, IHC, n=31) and BAP1 low (TCGA, RNAseq, n=40) The expression levels of *GPX4*, *SCL7A11* and *BAP1* were split at the median, and curves were plotted using SPSS. BAP1 levels were determined via pathological analysis (IHC) in the LUMC UM cohort and divided based on transcription levels along the median in the TCGA.

Prior to assessment of the efficacy of ferroptosis induction *in vivo*, we analyzed the relation between the three major eukaryotic ROS detoxifying enzymes, catalase (*CAT*), superoxide dismutase 2 (*SOD2*), and glutathione peroxidase 4 (*GPX4*) on metastasis development and survival in two cohorts of UM. We found that out of *CAT*, p=0.25, *SOD2,* p=0.83 and *GPX4* p=0.041, only *GPX4* expression negatively correlated with UM-specific survival in the TCGA cohort (80 cases (Figure S3). Analysis of *GPX4* expression levels and the relation between *GPX4* and overall survival in primary UM patients in the TCGA were analyzed using Gene Expression Profiling Interactive Analysis (GEPIA2)^43, 44^. We noted a significant reduction in melanoma-related death in patients expressing high levels of *GPX4* in a cohort of 64 UM from the LUMC as assessed by micro-array, and in the development of metastases in The Cancer Genome Atlas cohort (TCGA, p<0.04, n=78) analyzed through GEPIA2 as shown in supplementary Figure S2^44^.

We focused on ferroptosis and some of the known key mechanisms that play a role in either *GPX4* function or intracellular iron metabolism because expression of both *GPX4* and glutamate/cysteine antiporter (system Xc-SCL7a11, p<0.001) showed a strong negative correlation with patient survival (Figure 5A, B). Moreover, both GPX4 and SCL7A11 showed an enhanced negative correlation with survival in patients with a loss of BAP1 (BAP1-) (GPX4 in BAP1-IHC negative cases (p<0.02) and SLC7A11 in BAP1 low TCGA cases (Figure 5B) (p<0.02)). These findings suggest that levels of ferroptosis detoxicating enzymes correlate with worse prognosis and that the UM prognostic marker BAP1 loss is predictive of the role of ferroptosis related genes (GPX4 and SCL7A11) in UM progression.

We subsequently tested the dose-dependent ferroptosis-mediated killing of a selection of primary (MP46, MEL285) and metastatic (Omm1, MM28, and Xmm66) UM cell lines by well-established ferroptosis inducers 1S,3R-RSL3 (RSL3) and Erastin^19, 20, 42^. Strikingly, all tested cell lines responded to the induction of ferroptosis in a dose-dependent manner, but their overall response rate *in vitro* was weak (Figure S4C). The strongest response was noted in UM cell line MEL285, an atypical UM cell line not carrying any of the hallmark UM driver mutations but expressing a high level of pERK, which is indicative of upstream RAS-RAF-MEK hyperactivation found in cutaneous melanomas^45^. This cell line expressed low levels of *GPX4*, high levels of SCL7A11 and was killed completely by 200 nM Erastin, while it showed approximately 80% growth reduction with 200 nM RSL3. Both effects could be almost completely rescued by addition of ferroptosis inhibitors ferrostatin or liproxstatin (Figure S4C), indicating that oncogenic RAS activation in UM cells pre-disposes these cells to high ferroptosis susceptibility. Both ferrostatin and liproxstatin are thought to act through trapping free radicals, causal to the peroxidation of cell membranes^46^. Activation of pro-ferroptotic signaling in cell line MEL285 was completely blocked by the addition of ferroptosis inhibitors, ferrostatin and liproxstatin, indicating a canonical pERK dependent ferroptosis activation in this cell line. All other tested cell lines showed a dose-dependent response to the induction of ferroptosis, although these cell lines could be rescued only marginally by the addition of either liproxstatin or ferrostatin. These *in vitro* data suggest that while UM cells are broadly susceptible to ferroptosis induction, there is a pre-disposition to ferroptosis induction in pERK positive cells. Most UM samples which could only be rescued by ferrostatin or liproxstatin treatment showed strong ERK activation (suggestive of upstream oncogenic RAS hyperactivation), indicating a canonical ferroptosis susceptibility profile. Moreover, *GPX4* protein levels (Figure S4B) seem to be indicative of ferroptosis resistance *in vivo*, further enhanced by the pro-ferroptotic upstream activation of RAS-RAF-MEK-ERK, either through via GNAq/GNA_11_ PKC or indirectly through RASGPR3, as reported by Moore *et al*., 2018^47^.

To test the converse of this hypothesis, namely if the *a priori* inhibition of ferroptosis before engraftment could prevent the injected cells from dying in circulation. To this end, we treated Xmm66 and Omm2.3 with the ferroptosis inhibitors ferrostatin and liproxstatin prior to zebrafish engraftment (Figure 6B). Both inhibitors significantly enhanced UM cell survival in the circulation for Xmm66 (P<0.001 for both Liproxstatin and ferrostatin), up to 24 hours after systemic implantation into zebrafish larvae. Cell line Omm2.3 showed a marked, yet statistically insignificant, increase in cancer cell burden. These results indicate that ferroptosis plays a key role in the curbing of metastatic dissemination of UM.

**Figure 6.**
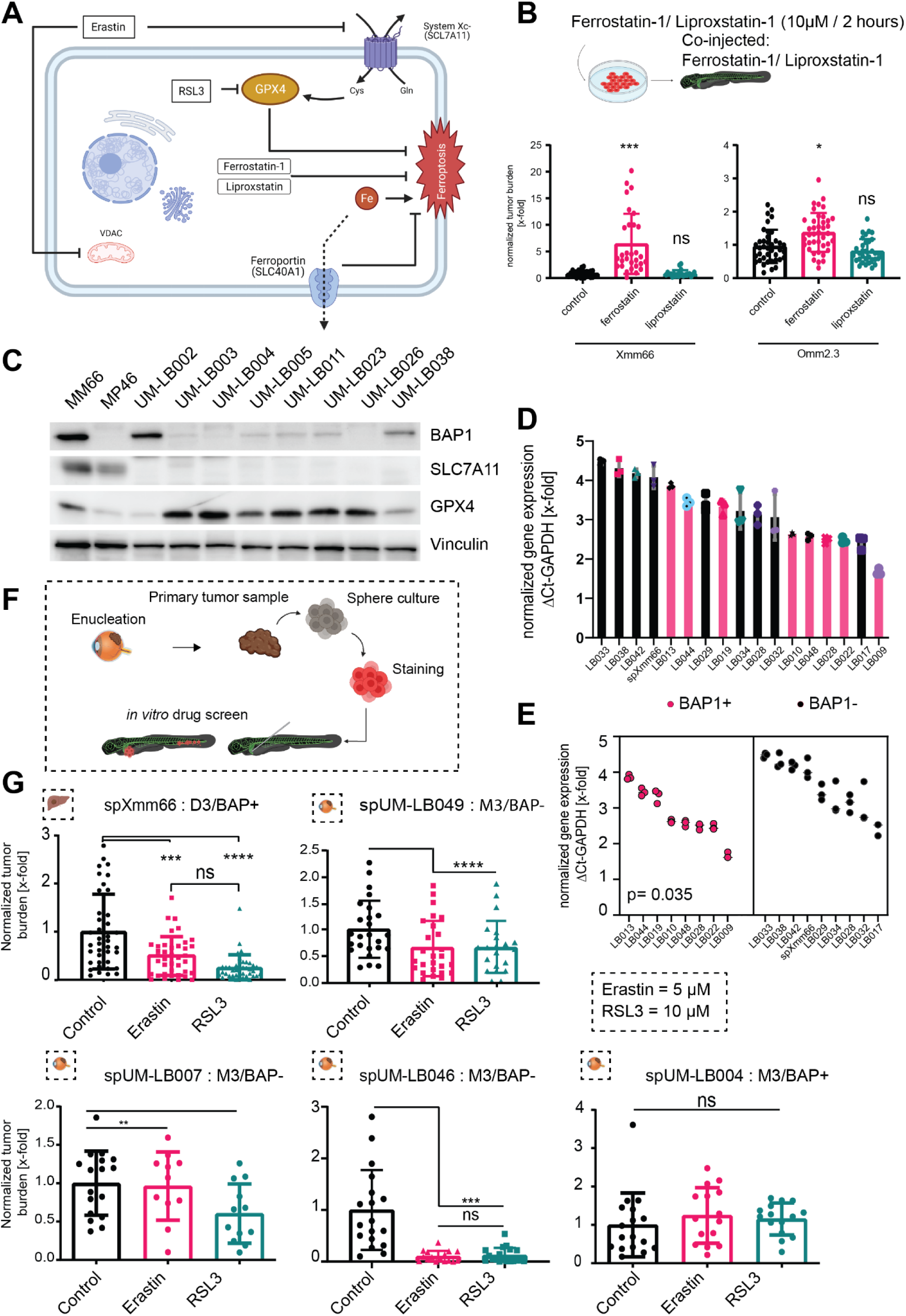
Treatment of UM patient-derived zf-Xenografts shows strong inhibitory potential of ferroptosis inducing compounds *in vivo*. A) Schematic representation of ferroptosis signaling. B) Pre-treatment of engrafted, adherent UM cell lines with anti-ferroptotic ROS traps ferrostatin-1 and Liproxstatin-1 prior to and its effect on UM cell survival *in vivo*. Whole body zebrafish measurements at 24 hours post injection, n=20, normalized to DMSO treated control, pre-treatment with ferrostatin-1 significantly enhances cell survival in cell line Xmm66 p<0.0003 and for Omm2.3 p<0.04 where in the liproxstatin pre-treated groups a slight enhancement of cell survival was measured (ns). Error bars represent SEM. C) Westernblot staining for *GPX4*, *SCL7A11* and BAP1, with vinculin as a loading control, showing high levels of SCL7A11 on both cell line samples MM66 and MP46 and undetectable levels on patient samples (UM-LB02-38), *GPX4* levels are elevated in patient samples in a BAP1-dependent manner. UM08003-26 are BAP1^-/-^ patient samples, low levels of BAP1 are detected in samples UM-LB003-26 most likely due to inclusion of stromal cells. D) qPCR analysis of GPX4 mRNA expression in primary UM tissues, with known BAP1 status, all samples measured in triplicate and normalized to GAPDH reference (ΔCT). E) Comparative analysis of GPX4 expression between BAP1+ and BAP1-patient samples indicating a significant enhancement in GPX4 levels in BAP1-patient samples. F) Schematical approach of zebrafish xenografts treatment of both primary and metastatic melanomas treated for 6 days with Erastin (5μM) or RSL3 (10μM) indicates a strong inhibition of normalized tumor burden. G) Zebrafish xenografts treatment of both primary and metastatic melanomas treated for 6 days with Erastin (5μM) or RSL3 (10μM), responding in a chromosome 3 monosomy/ Bap 1 dependent manner, with the exception of spXmm66 (D3/Bap 1^+^). Normalized tumor burden shown, n=40 (spXmm66) and n=20 (spUM-LB046, spUM-LB049), error bars show ±SEM.

To test the long-term influence of ferroptosis *in vivo*, we first established the MTD for both Erastin and RSL3 on un-injected zebrafish larvae: these were 5 µM and 10 µM, respectively (Supplementary Figure 2C). We subjected 36 engrafted zebrafish larvae to either compound, alongside DMSO as vehicle control, changing the compound-containing water or vehicle every other day (Schematically represented in Figure 6A). After 5 days of treatment, we imaged 20-40 whole zebrafish larvae per condition using a fluorescent stereo microscope, and quantified the red fluorescent integrated density, as a measure of cancer cell survival, normalizing to DMSO control (as described by Groenewoud *et al.* 2021)^29^.

In addition, we examined a panel of 8 primary UM patient samples, UM08002-UM08038 for the presence of *GPX4*, SCL7A11, and the BAP1 protein, to determine if there was a correlation between BAP1 expression and the ferroptosis-related proteins *GPX4* and *SCL7A11*. These samples were compared to established metastatic UM cell lines MM66 (*BAP1* positive) and MP46 (*BAP1* negative) (Figure 6C). We found that the two cell lines showed highly-elevated levels of SCL7A11 when compared to the patient-derived samples, in the case of MM66 in the absence of BAP1 mutation. The primary patient samples showed a positive correlation between *GPX4* levels and *BAP1* loss (UM08002 and UM08038 showed BAP1 expression and a low expression of *GPX4*). In parallel, we determined the dependence of GPX4 expression on BAP1 presence or absence. To this end, we performed a confirmatory qPCR-based analysis of GPX4 expression for two primary UM patient cohorts (BAP1^+^=8, BAP1^-^=9) and detected that GPX4 high and low populations could be segregated based on BAP1 status (p=0.035, Figure 6D, E). None of the primary samples exhibited detectable levels of SCL7A11 and hence no correlation with BAP1 levels could be determined. We examined the susceptibility of metastatic (spXmm66) and primary UM (spUM-LB046, spUM-LB049, spUM-LB007 were all BAP1^-^ and spUM-LB004 was BAP1^+^) to the induction of ferroptosis *in vivo,* monitored as reduction of relative tumor growth in zebrafish xenograft model (Figure 6G). Upon in vivo treatment, all but one primary UM samples (with the exception of BAP1 proficient sample spUM-LB004), as well as the metastasis-derived spXmm66 sample, displayed ferroptosis when challenged with RSL3 and Erastin (spUM-LB046 p<0.0001 both; spUM-LB049 ns and p=0.03; spUM-LB004, ns; spUM-LB007, p=0.03 and ns). Strikingly, although spXmm66 is derived from a BAP1^wt^ tumor it has high *GPX4* protein levels (data not shown), explaining its strong response to RSL3 and to a lesser extent Erastin (reduction of tumor burden, p<0.001 and <0.001, respectively).

In conclusion, using this zebrafish model, we have been able to demonstrate that both metastatic and primary UM cells were susceptible to pharmacological ferroptosis induction, manifesting stronger inhibition *in vivo* compared to *in vitro* (Figure 6G). Moreover, we have shown a possible predictor for ferroptosis treatment response in both clinically relevant and routinely detected UM markers BAP1 and monosomy 3, indicating that these patients could benefit from pro-ferroptotic therapy.

## Discussion

We have been able to establish a robust protocol for the generation of metastatic UM spheroid cultures. Using this method, we generated one continuous UM culture of metastatic origin and several short-lived primary and metastatic spheroid cultures. In concordance with the general disease progression of UM and the lack of available metastatic UM material, we chose to focus on primary UM samples. In choosing these, we reasoned that our screening efforts could possibly be translated to assumed metastatic patients, providing data for adjuvant treatment. We combined our model with the knowledge of the strong prognostic value of monosomy 3/BAP1 loss to pre-screen patients with a high risk of developing metastatic UM. We developed our near patient zf-PDX model using the highly proliferative spheroid culture spXmm66, seeing as it is the only biological UM entity that readily yields perivascular metastatic colonies (Figure 3B). In concordance with these findings, we were incapable of robustly generating metastatic colonies after injection of stable adherent cell lines Omm1, Omm2.3, Omm2.5 and Xmm66 (data for Omm1, Omm2.5 not shown). Furthermore, our data indicate that the metastatic capacity of spheroid line spXmm66 is dependent on an absence of ROCK signaling. When spXmm66 was cultured in adherent culture for one week, this completely abrogated this cell line’s intrinsic metastatic capacity. Inclusion of ROCK inhibitor Y267632 during the adherent culture of spXmm66 significantly reduced the loss of metastatic capacity, whereas inclusion of serum enhanced the loss of metastatic capacity. This leads us to conclude that for spXmm66, its metastatic potential is linked to its spheroidal low RhoA signaling nature; we assume that there is a mechanical force-dependent mechanism responsible for the loss of metastatic potential, which is further potentiated by the addition of serum to the medium. To exclude a direct effect of the stem cell medium, we cultured Xmm66 and Omm2.3 in complete NSC medium, but observed no enhancement of metastatic potential, neither in the adherent nor the spheroidal culture.

Subsequently we validated the zf-PDX model through single agent and combinatorial drug treatments, using experimental treatments developed to inhibit both cell line Xmm66 and murine localized tumor model. Metastatic UM PDX model spXmm66 largely recapitulated the findings in murine models, indicating that it could be used to assess drug efficacies *in vivo* (Figure 7).

**Figure 7.**
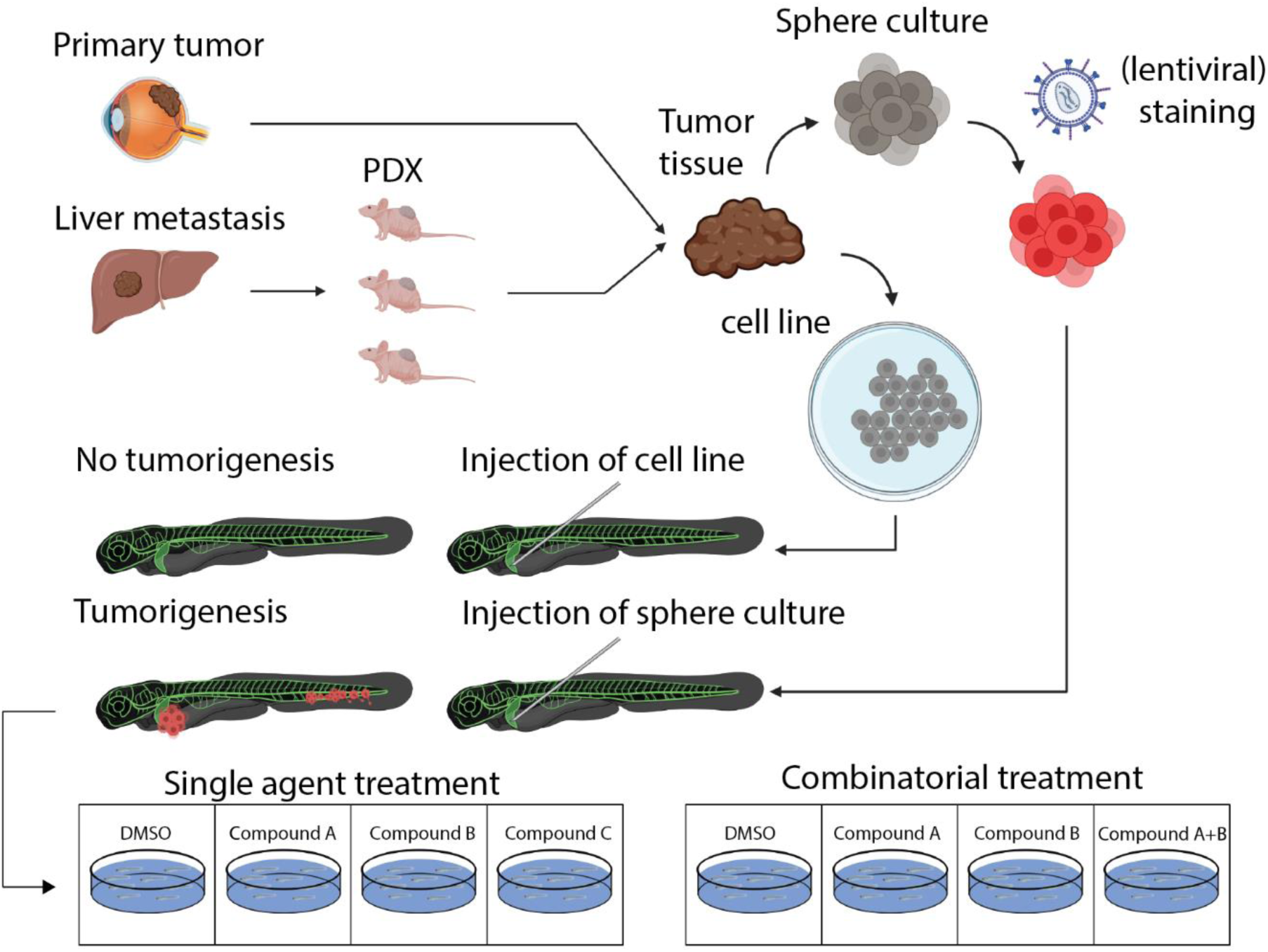
Establishment of a zebrafish uveal melanoma PDX model. A) Metastatic (or primary) UM are collected and are used to establish mouse xenografts and effectively propagated in NOD-SCID mice (via sub-cutaneious engraftment) or are directly used to establish a non-adherent near-patient spheroid culture (as shown under B). C) The establishment of these spheroid cultures allows for the *in vitro* (lentiviral) modification of patient material (addition of molecular tracers, reporters, etc.) prior to engraftment, allowing the separation of one biological sample over two individual experiments. In some cases, the establishment of spheroid culture allows for the generation of long-lived (p>20) metastatic UM lines, allowing the in-depth analysis of metastatic UM and drug screening. Spheroids are dissociated prior to engraftment (either physically or enzymatically) and the single cell suspension derived thereof is engrafted through the Duct of Cuvier (the embryonic common cardinal vein) of 2dpf zebrafish larvae, approximately 250-350 cells per larva. Zebrafish are screened 1 dpi, where all the selected, positively engrafted larvae, are randomly divided into groups treated with either vehicle, compound A, compound B or a combination of both compound A and B (all at the respective maximum tolerated dose of the combination of compound A and B). Anti-tumor efficacy of all groups is determined through an integrated density measure using FIJI, based on standardized fluorescent micrographs. All measures are subsequently normalized to vehicle control and shown as normalized tumor burden.

Taking together the overall lack of tumorigenic capacity of adherent UM cells and the rapid clearance of UM cells after hematogenous engraftment we reasoned that there was a cell intrinsic mechanism responsible for the obvious disconnect between the high metastatic capacity of patient derived cells and *in vitro* propagated UM cells. Reactive oxygen and more specifically ferroptosis has recently been linked to the curbing of metastatic spread of cancer cells, specifically acting on circulating cancer cells.

To assess the efficacy of ferroptosis induction on UM cells *in vivo,* we selected two distinct ferroptosis inducers, Erastin and 1S, 3R-RSL3 (RSL3), which are, respectively, class I and class II ferroptosis inducers (FINs). Erastin inhibits the glutathione/cysteine antiporter function of system Xc- (encoded by SLC7a11 and SLC3a2), undermining GPX4’s capacity to catalyze phospholipid peroxidation by depriving it of its substrate^48^. The pro-ferroptotic effect of Erastin is further enhanced by its action upon mitochondrial voltage-dependent anion channel 2 (VDAC2). VDAC2 inhibition leads to a massive increase in intracellular ROS levels through a disruption of the mitochondrial membrane potential, followed by permeation of ROS into the cytosol proper^40^. Interestingly, we found that induction of ferroptosis is highly effective against spreading of primary UM cells *in vivo* (Figure 6G), and shows a strong inhibitory tendency for all BAP1- samples tested. Furthermore, the recent publication of Luo & Ma, 2021 underscores the presence of a ferroptosis-related gene signature among UM tumors ^49^. This finding, taken together with the TCGA and LUMC patient cohort-derived data, confirms the presence of a strong negative correlation of the ferroptosis-related genes *GPX4* and system Xc- (*SCL7A11*) with and (metastasis-free) survival. Furthermore, our zf-PDX based near-patient *in vivo* drug screen indicates the translational value and validity of ferroptosis inducers for the treatment of UM.

Subsequent *in vitro* data showed that most cultured UM cells are largely refractory towards ferroptosis induction at similar concentrations (Erastin 5 µM and RSL3 10 µM *in vivo*) as used during successful *in vivo* ferroptosis induction. Taken together, our findings underscore the validity of using the zebrafish model for the discovery of novel cancer therapeutics, in this instance for a malignancy where until now there was no available metastatic animal model. Ultimately, our findings add to the building body of evidence supporting the value of ferroptosis induction as a potential treatment in metastatic uveal melanoma^50^.

## Author’s contribution

A.G. designed, performed and analyzed all *in vivo* experiments, performed GEPIA analysis, wrote the manuscript and designed the graphics. J.Y. performed additional zebrafish engraftments, Western blots and qPCR. M.C.G. provided information on the patients and performed the statistical analysis on patient information, S.A., F.N. and D.D. supplied the metastatic UM material, H.K. and S.C. performed the immuno-stains and the analysis thereof, A.G.J. performed the *in vitro* proliferation assays and Western blots and provided assessment of this manuscript. M.J. provided primary UM tissue samples, patient data and assessment of this manuscript, E.S.J. provided funding and supervised the project.

## Acknowledgements

We kindly thank Emilie Vinolo for the managerial assistance provided during this UMcure2020 project.

We further thank Amina Teunisse for technical assistance.

This work has received funding from the European Union’s Horizon 2020 research and innovation program under grant agreement No 667787 (UM Cure 2020 project, www.umcure2020.org

All graphics (excluding scientific data) were generated using Biorender.com

The results shown here are in whole or part based upon data generated by the TCGA Research Network: https://www.cancer.gov/tcga

