## supplementary figure 2 for "Patient-derived zebrafish xenograft models reveal ferroptosis as a fatal and druggable weakness in metastatic uveal melanoma"

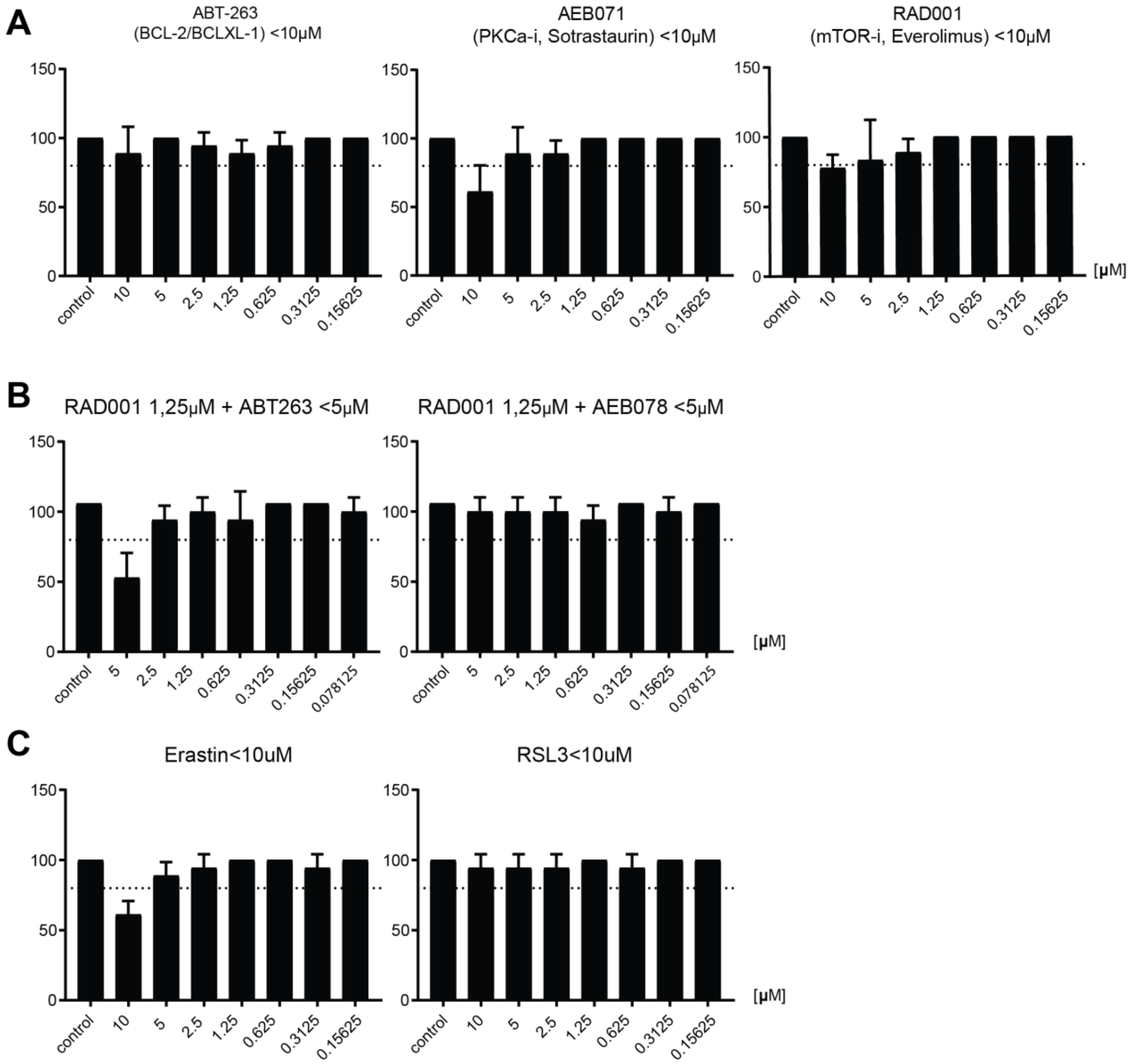


**Supplementary Figure 2. Establishment of maximum tolerated dosage of tested putative anti-UM therapeutics.** Dose response graphs on treated, uninjected zebrafish. Treatment concentration were deemed to be viable when at least 80% of the treated, uninjected, embryos survived for the duration of the treatment. A) First mono treatments were established (ABT-263, AB071 and RAD001) B) whereafter the combinatorial treatments were established (RAD001+ABT-263 and RAD001+AEB071). C) Ferroptosis inducing compounds. All treatments were refreshed every other day, for 5 days post injection, up to the final day of the experiment (8 days post fertilization).
