## supplementary figure 3 for "Patient-derived zebrafish xenograft models reveal ferroptosis as a fatal and druggable weakness in metastatic uveal melanoma"

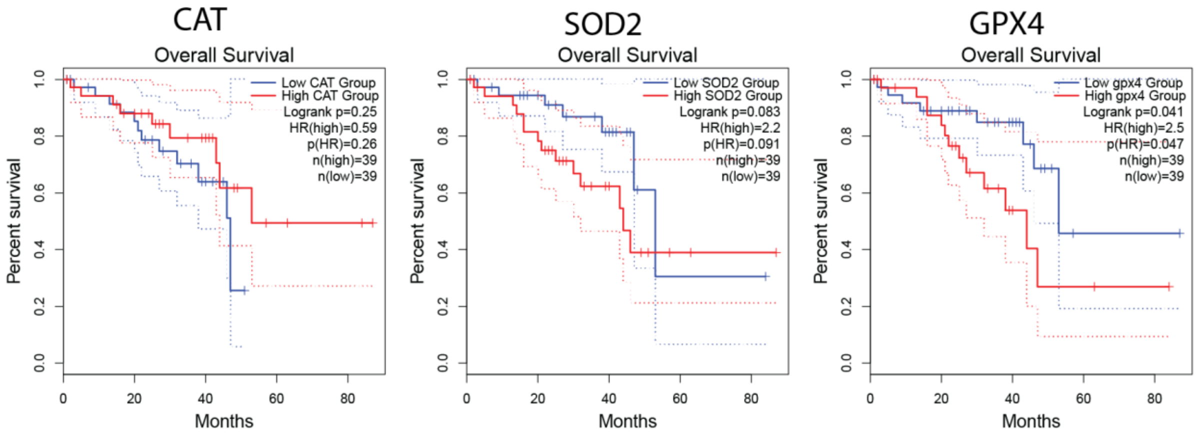


**Supplementary Figure 3. In silico analysis of general ROS detoxifying enzymes in UM indicates that ferroptosis-related genes are strongly associated with a bad prognosis in UM.** Analysis of the cancer genome atlas (TCGA) revealed that of the three major ROS detoxifying enzymes catalase (CAT), superoxide dismutase2 (SOD2) and glutathione peroxide 4 (*GPX4*), *GPX4* is the only one that correlates significantly with a bad prognosis.
