## supplementary figure 4 for "Patient-derived zebrafish xenograft models reveal ferroptosis as a fatal and druggable weakness in metastatic uveal melanoma"

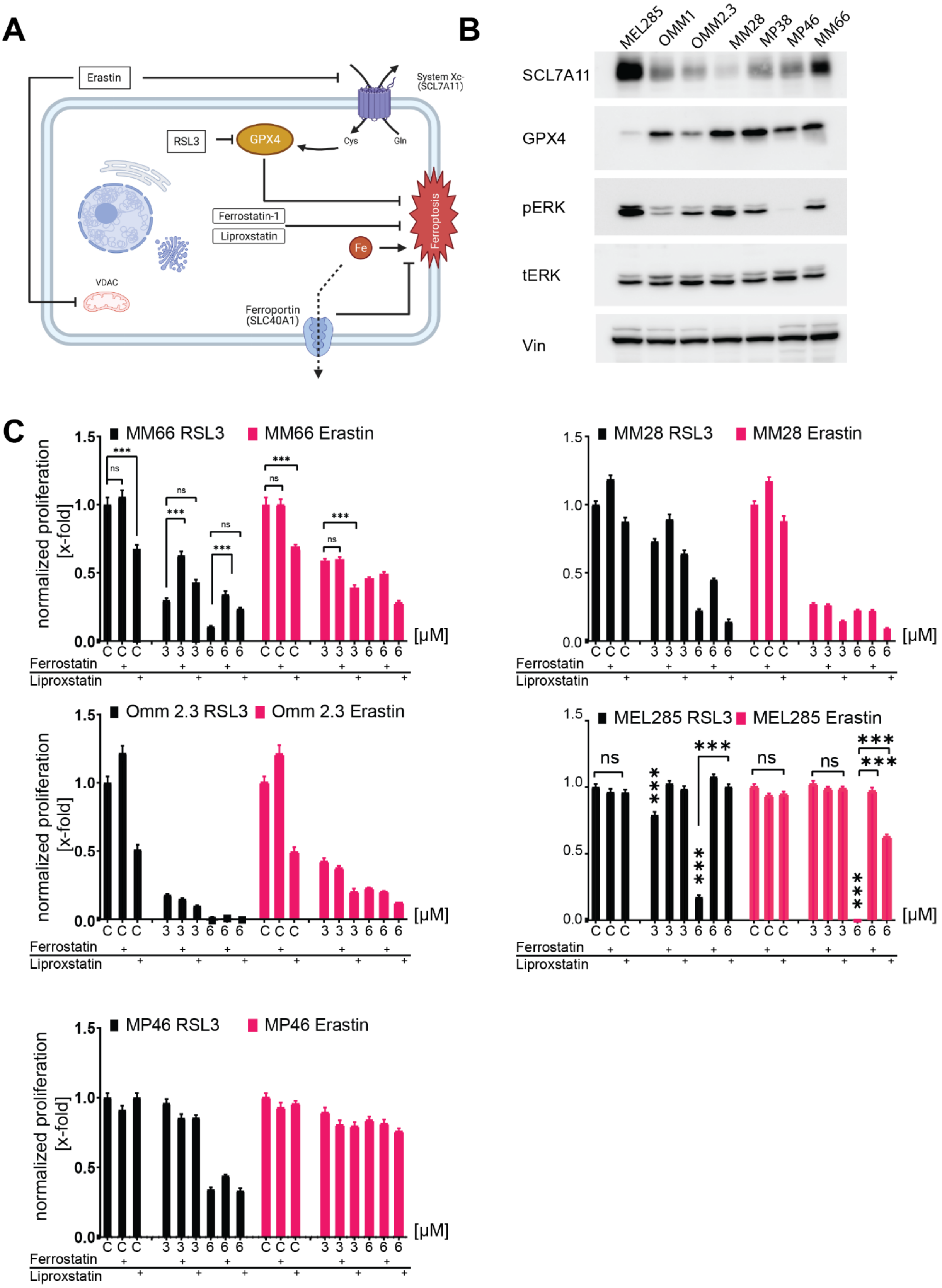
A
Erastin
System Xc-
(SCL7A11)
RSL3
GPX4
Cys
Gln
Ferrostatin-1
Liproxstatin
Ferroptosis
Fe
VDAC
Ferroportin
(SLC40A1)
B
MEL285
OMM1
OMM2.3
MM28
MP38
MP46
MM66
SCL7A11
GPX4
pERK
tERK
Vin
C
MM66 RSL3
MM66 Erastin
MM28 RSL3
MM28 Erastin
Omm 2.3 RSL3
Omm 2.3 Erastin
MEL285 RSL3
MEL285 Erastin
MP46 RSL3
MP46 Erastin
normalized proliferation [x-fold]
[µM]
Ferrostatin
Liproxstatin

**Supplementary Figure S4**. Induction of ferroptosis significantly reduces cell survival *in vitro*. A) Induction of ferroptosis through inhibition of system $X_c$ (Erastin, ER) or through inhibition of glutathione peroxidase 4 (*GPX4*) with RSL3 shows effective reduction of viability of both primary and metastatic uveal melanoma cells *in vitro* A) *in vitro* treatment of primary (MP46) and metastatic (Omm1, mm28 and Xmm66) uveal melanoma.  B) Westernblots detecting GPX4 and System Xc- (SCL7A11) and concordant expression of total ERK (tERK) and phosphorylated ERK (pERK) indicative of upstream RAS activation. C) Rescue experiment, All cell lines were treated with 8 and 4 µM Erastin and 6 and 3 µM RSL3, with the exception of MEL285 which was treated with 0.2 and 0.05 µM Erastin or RSL3 and subsequent rescue was attempted with ferroptosis inhibitors ferrostatin and Liproxstatin.
