## supplementary figure 5 for "Patient-derived zebrafish xenograft models reveal ferroptosis as a fatal and druggable weakness in metastatic uveal melanoma"

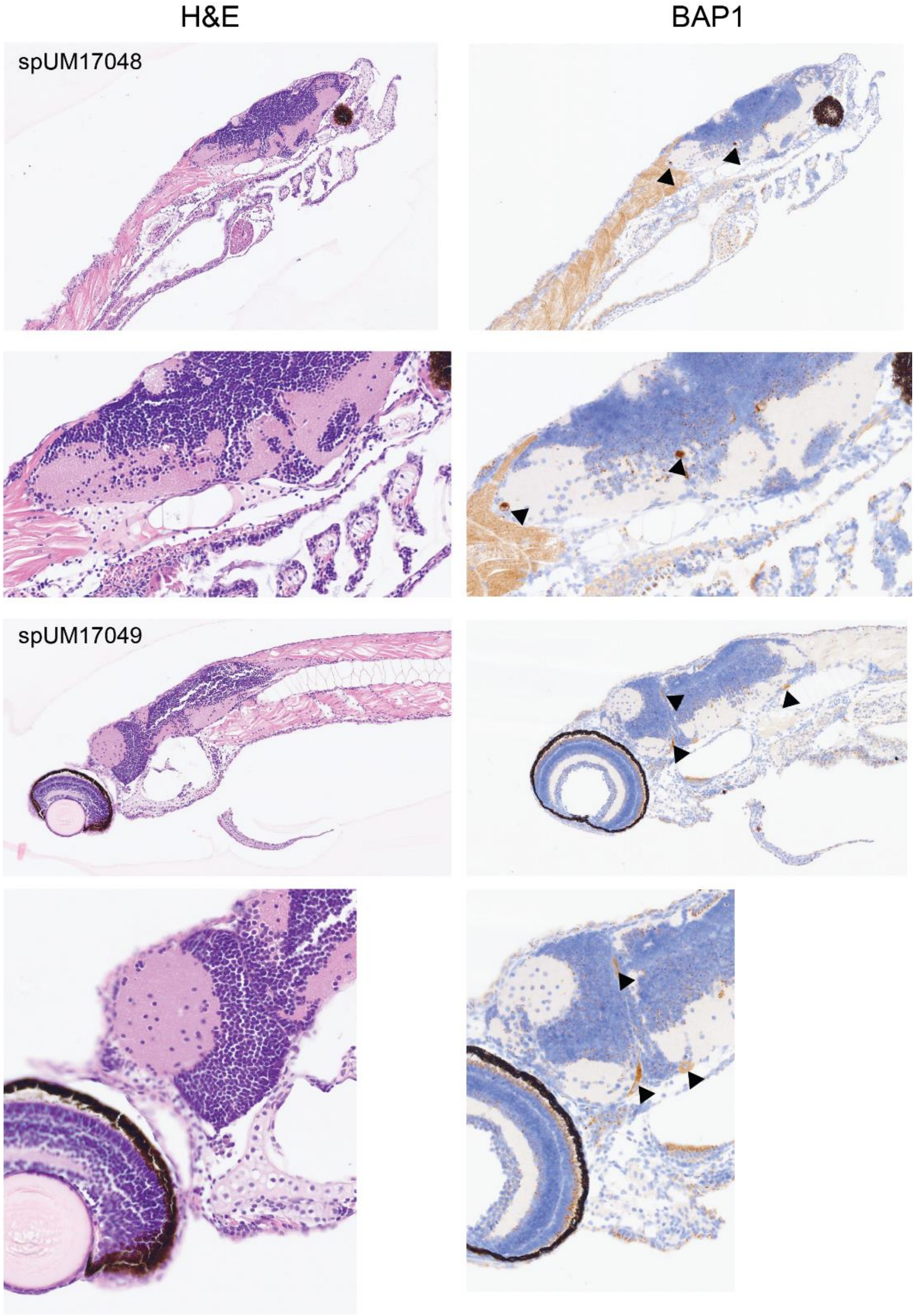


**Supplementary Figure S5. Additional IHC staining of engrafted primary UM samples.** Zebrafish larvae injected at 48 hpf, fixed at 6 dpi, fixed and oriented in a low melting temperature agarose block. Presence of BAP1 was assessed as previously described.
