## supplementary figure 1 for "Patient-derived zebrafish xenograft models reveal ferroptosis as a fatal and druggable weakness in metastatic uveal melanoma"

### Supplementary figures

-

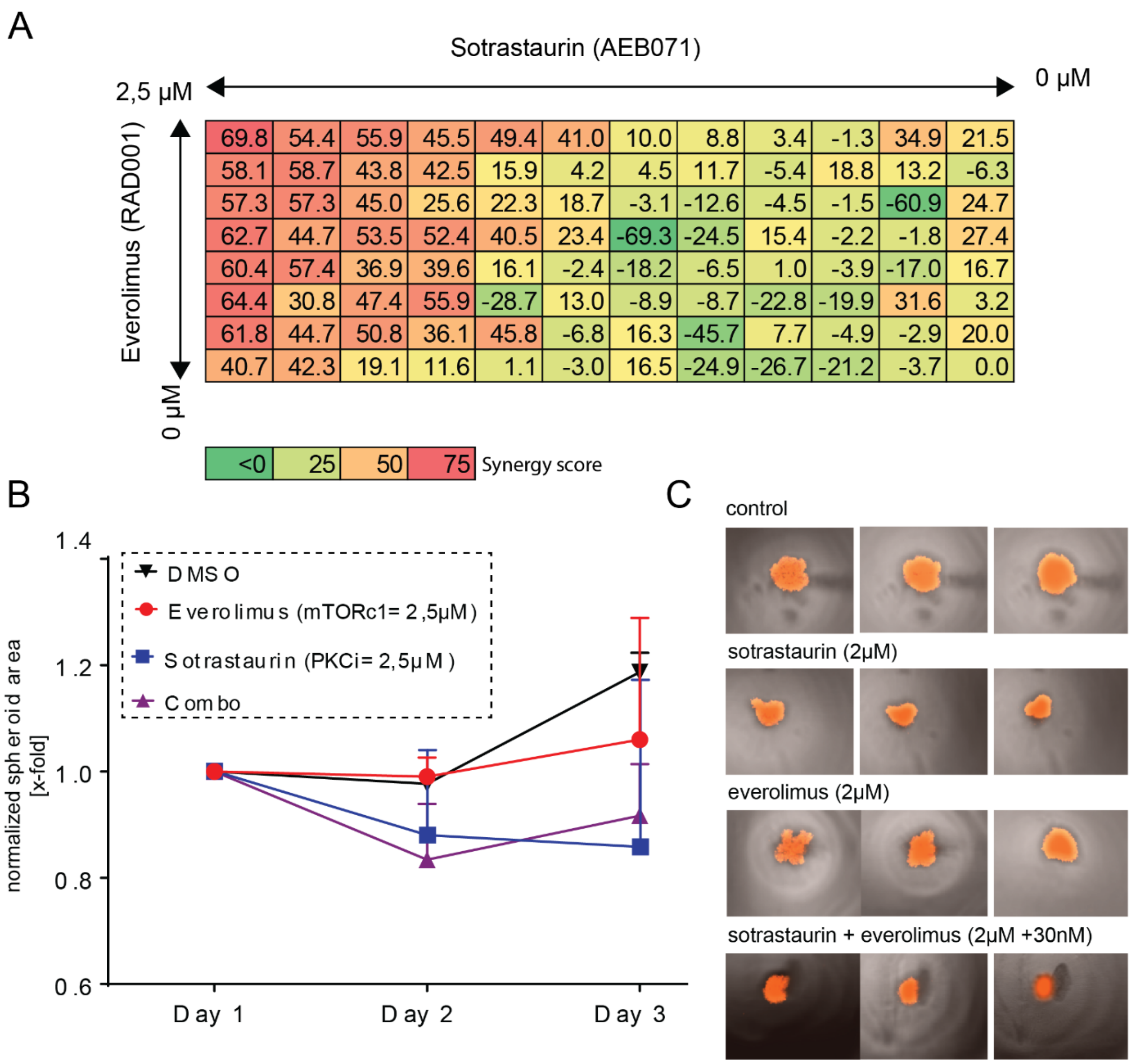


**Supplementary Figure 1. Spheroid culture-based drug screening method for pre-clinical evaluation of combinatorial drug treatment.** A) Heatmap of drug synergy between PKC inhibitor sotrastaurin and mTORC1 inhibitor everolimus, tested on spheroid culture line spXmm66, inhibition indicated in percentages and shown graphically as a heat-map (green to red, antagonistic to synergistic, respectively) made as an end point measurement after 3 days of treatment, using cell CellTiter Glo 2.0 as per the manufacturer's prescription. B) cell growth kinetics measured over time (based on spheroid surface area), measured on 1-,2- and 3-days post seeding in ultra-low adhesion 96-wells plates. C) Fluorescent micrographs of spXmm66 spheroids after 3 days of treatment with either sotrastaurin 2 µM, everolimus 2 µM the combination of both (sotrastaurin 2 µM and everolimus at 30 nM) compared to vehicle control (DMSO).
