## supplementary table 1 for "Patient-derived zebrafish xenograft models reveal ferroptosis as a fatal and druggable weakness in metastatic uveal melanoma"

Supplementary Table ST1 Overview of tissues used for spheroid culture derivation

| | Name | Recovered from freezing | Sphere culture established | Lentiviral transduction | zebrafish xenograft | Validation | Growth speed | Maximum passage number | Reference(s) |
|---|---|---|---|---|---|---|---|---|---|
| Metastatic | MM26 | ✓ | ✓ | - | ✓ | S, IHC, C | -/± | 4 | Nemati, Laurent |
| | MM28 | +/- | +/- | - | IM | N/A | - | 4 | Nemati, Amirouchene |
| | MM33 | ✓ | ✓ | - | IM | S | -/± | 4 | Nemati, Laurent, Carita |
| | MM52 | ✓ | ✓ | - | IM | N/A | -- | 4 | Nemati, Laurent, Carita |
| | MM66 | ✓ | ✓ | ✓ | ✓ | S, IHC, C, D | + | 20+ | Nemati, Laurent, Amirouchene |
| | MM252 | N/A | ✓ | - | IM | S | N/A | N/A | N/A |
| | MM257 | N/A | ✓ | - | IM | S | N/A | N/A | N/A |
| | MM267 | ✓ | ✓ | ✓ | IM | S | N/A | N/A | N/A |
| | MM278 | N/A | ✓ | - | IM | S | N/A | N/A | N/A |
| | MM293 | ✓ | ✓ | ✓ | IM | S | N/A | N/A | N/A |
| | MM299 | N/A | ✓ | - | IM | S | N/A | N/A | N/A |
| | MM300 | N/A | ✓ | ± | IM | S | N/A | N/A | N/A |
| | MM309 | ✓ | ✓ | ✓ | IM | S | N/A | N/A | N/A |
| | MM325 | ✓ | ✓ | ✓ | IM | S | N/A | N/A | N/A |
| Primary | UM 17-045 | ✓ | ✓ | N/A | - | S | N/A | N/A | N/A |
| | UM 17-046 | ✓ | ✓ | ✓ | ✓ | S, D | N/A | N/A | N/A |
| | UM 17-047 | ✓ | ✓ | N/A | ✓ | S, C | N/A | N/A | N/A |
| | UM 17-048 | ✓ | ✓ | N/A | ✓ | S, C, D | N/A | N/A | N/A |
| | UM 17-049 | ✓ | ✓ | N/A | - | S | N/A | N/A | N/A |
| | UM 18-004 | ✓ | ✓ | N/A | ✓ | S, D | N/A | N/A | N/A |
| | UM 18-005 | ✓ | ✓ | N/A | - | S | N/A | N/A | N/A |
| | UM 18-007 | ✓ | ✓ | N/A | ✓ | S, D | N/A | N/A | N/A |
| | UM 18-008 | ✓ | ✓ | N/A | - | S | N/A | N/A | N/A |
| | UM 18-010 | ✓ | ✓ | N/A | - | S | N/A | N/A | N/A |

S=Sphere culture, IHC=Immunohistohemistry, C=confocal imaging, D=Drug screen
IM= insufficient material
