## supplementary table 2 for "Patient-derived zebrafish xenograft models reveal ferroptosis as a fatal and druggable weakness in metastatic uveal melanoma"

Supplementary table ST2 qPCR primer sequences

| | FW | | |
|---|---|---|---|
| MITF | 5'- | AACAGAGAGTGCCCGTGAGT | -3' |
| TYR | 5'- | TACGGCGTAATCCTGGAAAC | -3' |
| DCT | 5'- | GGGAGGAACGAGTGTGATGT | -3' |
| TYRP1 | 5'- | CTGGAATTTTGCAACGGGGA | -3' |
| SOX10 | 5'- | CTTCATGGTGTGGGCTCAG | -3' |
| GPX4 | 5'- | TGGACAAGTACCGGGGCTTC | -3' |
| SCL7A11 | 5'- | TGCTGTGATATCCCTGGCAT | -3' |

| | RV | | |
|---|---|---|---|
| MITF | 5'- | GACATGGCAAGCTCAGGACT | -3' |
| TYR | 5'- | ATTGTGCATGCTGCTTTGAG | -3' |
| DCT | 5'- | TGGCAATTTCATGCTGTTTC | -3' |
| TYRP1 | 5'- | CCATCCTCGGTGCTGTTACA | -3' |
| SOX10 | 5'- | TGTAGTCCGGGTGGTCTTTC | -3' |
| GPX4 | 5'- | CGAACTGGTTACACGGGAAG | -3' |
| SCL7A11 | 5'- | AGCTGCATAACTCCAGGGAC | -3' |
